# Recovering Hidden Neural Oscillatory Modes from Partial Observations through Delay Embedding and Perturbation

**DOI:** 10.64898/2026.10.05.756919

**Authors:** Raku Miyoshi, Mikito Ogino, Daiki Sekizawa, Masafumi Oizumi

## Abstract

Characterising the oscillatory behaviour generated by neural interactions is central to systems neuroscience because these dynamical properties are linked to brain function. However, inferring latent neural dynamics is challenging because recordings sample only part of the system. Time-delay embedding allows dynamical modes to be estimated from partial recordings, using methods such as Hankel dynamic mode decomposition (Hankel DMD). Despite its promise, two questions remain: what dynamical information is recoverable through time-delay embedding, and how can recovery be improved? We address both questions using control theory. Our framework centres on factorising the Hankel matrix of delay-embedded observations into an observability matrix and a trajectory matrix with the form of a controllability matrix. We show that their ranks impose complementary constraints on what any delay-embedding method can recover: delay embedding can increase observability rank, while perturbation can increase trajectory rank by exciting otherwise inactive modes. We validate this framework using Hankel DMD in simulations and human electrocorticography (ECoG). In both, perturbation increased the number of fast-decaying mode estimates, whereas greater delay depth increased the number of reliably estimated modes and, in ECoG, extended their frequency range. These results clarify what partial recordings can reveal about neural dynamics and how to reveal more.

## 1 Introduction

Interactions among neuronal populations generate collective dynamics that support neural communication and information processing in the brain. Characterising the oscillatory organisation of these dynamics is important for understanding sensory processing, attention, and memory [1, 2]. For example, selective visual attention modulates neuronal synchronisation, while experimentally induced gamma rhythms alter sensory responses [3, 4]. Neural oscillations also exhibit distinct spectral patterns across sleep and wakefulness and with changes in arousal, accompanying changes in how neural circuits process information [5–9]. Characterising the frequencies and decay rates of neural modes is therefore important for understanding the temporal organisation of neural dynamics.

Dynamic mode decomposition (DMD) provides a data-driven approach to characterising these oscillatory dynamics by extracting spatiotemporal modes from multivariate time series [10–12]. Each estimated mode combines a spatial activity pattern with a temporal frequency and a growth or decay rate, describing where activity is expressed, how rapidly it oscillates, and how it evolves over time. Originally developed in fluid dynamics, DMD has since been applied to neuroscience, alongside related Koopman methods, to characterise electrophysiological network dynamics [13–16], identify dynamical patterns and predict behavioural differences from fMRI [17, 18], analyse whole-brain dynamics and arousal-related activity in calcium recordings [19, 20], and compare neural population dynamics across tasks and recording sessions [21, 22]. These methods offer a way to characterise the dynamical organisation of neural activity through the modes expressed in recorded time series.

However, accurately estimating the oscillatory modes of the underlying neural system from recordings faces two fundamental limitations. First, neural recordings provide only partial access to the underlying dynamical system. Whether obtained using EEG, ECoG, multielectrode recordings, Neuropixels probes or optical imaging, measurements sample only a subset or a coarse-grained representation of the latent neural state [23–25]. Consequently, distinct latent modes may be indistinguishable from instantaneous measurements. Second, even a mode that is observable in principle can be difficult to identify if it is weakly represented within the recording window. For example, during spontaneous activity, the initial condition and endogenous inputs may contain little contribution from particular modes, and rapidly decaying modes may decay before the observation window begins or fall below the noise level [26]. Modes absent from the recorded activity cannot be recovered from those recordings alone, while weak contributions make reliable estimation difficult. These limitations raise two linked questions: how do observation and excitation jointly constrain spectral recovery, and how can each constraint be alleviated?

To address these difficulties, we develop a framework based on control theory to clarify what dynamical information can be recovered from partial observations and how recovery can be improved. Specifically, we use the factorisation of the Hankel matrix, which combines observations with their time-delayed copies. For a linear system, this matrix factorises into two components: one describing how latent states appear in the delayed measurements and another describing which dynamical directions are represented in the recorded trajectory. These are the observability matrix and the trajectory matrix, respectively; the latter takes the form of a controllability matrix [27–30]. The ranks of these factors impose complementary constraints on the recoverable spectrum and clarify how two strategies can improve recovery. Delay embedding uses the temporal evolution of measurements to distinguish latent components that are indistinguishable at a single time point [31, 32], potentially increasing the observability rank. Perturbation can increase the trajectory rank by exciting modes that are otherwise inactive, or strengthen weak modal contributions and make them easier to estimate [26, 33]. Complete recovery of the latent spectrum in linear systems therefore requires both sufficient observability and sufficient excitation.

We first validate the theoretical framework in simulations and then examine its implications in human ECoG recordings. In a partially observed linear system with known eigenvalues, we independently vary the delay-embedding dimension and perturbation. Complete eigenvalue recovery occurs only when delay embedding and perturbation are combined, revealing modes that remain unidentified when either strategy is used alone. We then examine how these strategies affect mode estimates in finite, noisy neural recordings using human ECoG data acquired during repeated single-pulse electrical stimulation [34]. We compare DMD estimates from pre-stimulus spontaneous activity and post-stimulus transient responses across delay-embedding dimensions and assess their reproducibility across independent trials. Electrical stimulation increases the number of rapidly decaying mode estimates, while delay embedding increases the number of reproducible estimates and extends their frequency range. Together, these results demonstrate how delay embedding and perturbation can improve the dynamical information extracted from partial neural recordings.

## 2 Theoretical framework

In this paper, we develop a theoretical framework to quantify the limits of reconstructing neural dynamics from partial observations and identify ways to increase the dynamical information that can be recovered. Specifically, we address two questions: (1) Which aspects of the dynamics of the full system can be recovered from recordings that capture only part of its activity, and what limits their recovery? (2) How can we improve recovery within these limits through recording, analysis, and perturbation?

Our framework addresses these two questions by drawing on observability and controllability, two concepts from control theory. Starting with a linear system, we first ask whether a sequence of partial observations can distinguish the latent state (Fig. 1A). We then collect such sequences into a Hankel matrix and factorise it into an observability matrix *P* and a trajectory matrix *Q* (Fig. 1D), whose connection to controllability we explain below. Together, these factors determine how many independent state directions are represented in the recording. We next establish when Hankel dynamic mode decomposition (Hankel DMD) recovers the underlying dynamical spectrum, before explaining how delay embedding (Fig. 1B), recording, and perturbation (Fig. 1C) can increase recovery. Finally, we extend these principles to nonlinear systems through Koopman operator theory and state the conditions under which a finite-dimensional representation supports the same spectral recovery argument.

**Figure 1.**
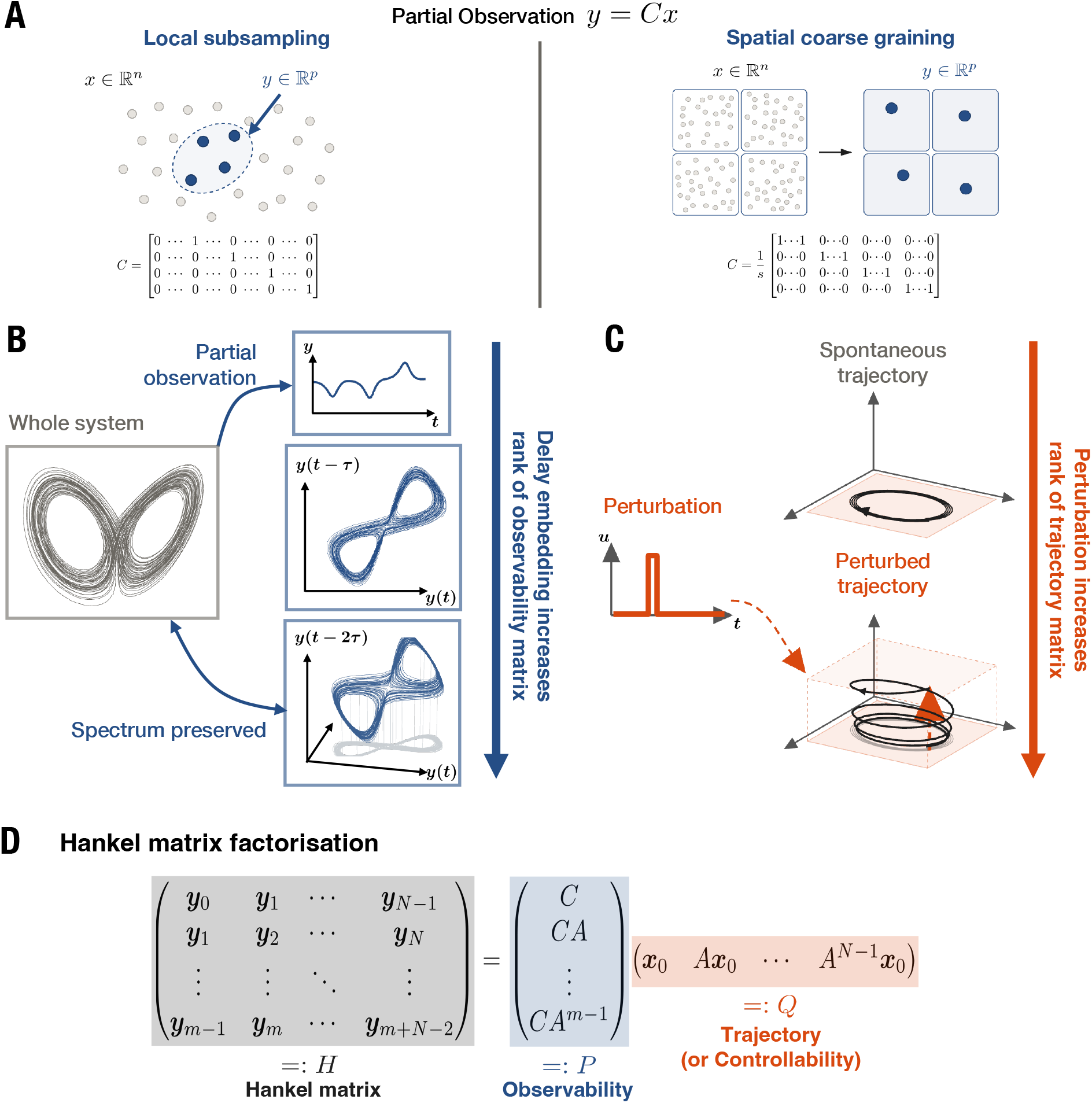
Delay embedding and perturbation reveal latent dynamics in the Hankel matrix. **(A)** Partial observations *y* = *Cx* obtained by local subsampling or spatial coarse graining. **(B)** Delay embedding increases the rank of the observability matrix *P* . **(C)** Perturbation excites additional dynamical directions, increasing the rank of the controllability matrix *Q*. **(D)** The Hankel matrix factorises as *H* = *PQ*.

### 2.1 Partial observation and the dynamics represented in a recording

Here we formalise the problem of reconstructing the full system’s dynamics from partial observation.

We first describe the full system by the linear evolution equation

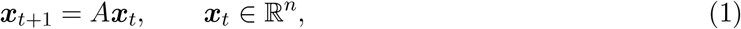

where ***x***_*t*_ is the underlying, or latent, neural state at time step *t*, including variables that are not directly measured. The matrix *A ∈* ℝ^*n×n*^ determines how this state evolves from one time step to the next. The recorded neural activity is given by

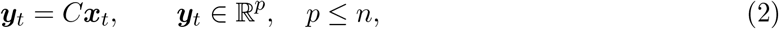

where the observation matrix *C ∈* ℝ^*p×n*^ specifies how the recording channels sample or combine the state variables (Fig. 1A). For example, for a single recording channel, *C* = [0, 1, 0, …, 0] selects the second state variable, whereas 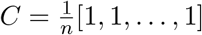 measures the average across all state variables.

The eigenvalues of *A* determine whether each dynamical mode grows, decays, or oscillates over time. For simplicity, suppose that *A* has *n* linearly independent eigenvectors. The state can then be expressed as a sum of modes,

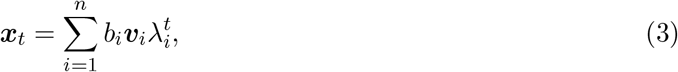

where ***v***_*i*_ *∈* ℂ^*n*^ is the *i*-th eigenvector of *A, λ*_*i*_ *∈* ℂ is its associated eigenvalue, and *b*_*i*_ *∈* ℂ is the initial contribution of that mode. Each eigenvector describes a pattern of activity across the state variables. Its eigenvalue determines how that pattern changes over time through the factor 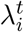. The collection of eigenvalues is called the dynamical spectrum. The magnitude and phase of an eigenvalue determine a mode’s growth or decay and oscillation frequency.

Our objective is to recover the dynamical spectrum, which is preserved under an invertible linear transform of state coordinates even though the entries of *A* change. Thus, even when the system states are only partially observed, we may be able to reconstruct the dynamics in a different set of coordinates with the same eigenvalues. Recovering the entries of *A* requires additional information about how the reconstructed coordinates relate to the original state variables; this information is usually unavailable. Given this limitation, the question to ask here is whether the recordings contain enough information to recover the underlying dynamical spectrum.

Time-delay embedding provides a route to reconstructing the underlying dynamical spectrum by combining measurements across time. Takens’ embedding theorem gives conditions under which delayed measurements can represent the underlying state and its dynamics in a different set of coordinates [31]. A delay embedding does not by itself guarantee a finite-dimensional linear representation; the conditions for spectral recovery are developed below. Inspired by Takens’ theorem, a variant of DMD called Hankel DMD was proposed to use these delayed measurements to estimate a linear model of how activity changes over time and extract its dynamical spectrum [32]. In the following subsections, based on Hankel DMD and control theory, we use observability and controllability to explain when this approach can recover the latent dynamics and how delay embedding and perturbation can improve recovery.

### 2.2 Limits of extractable information from partial observation: Hankel matrix factorisation

To understand how much information partial recordings provide about the latent dynamics, consider how the observed activity changes over time. A single measurement may not distinguish different latent states, but their subsequent measurements may differ as unobserved components influence the recorded variables (Fig. 1B). We therefore consider a sequence of successive measurements together.

We consider the recordings of the autonomous linear system in Eqs. (1) and (2), with fixed *A* and *C*. Stacking *m* successive measurements gives

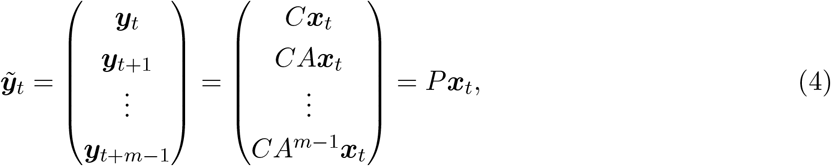

where

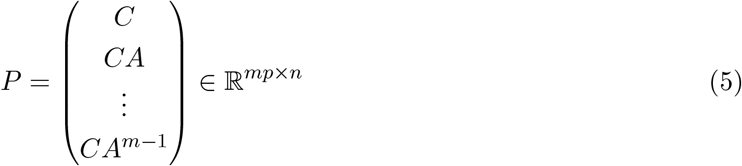

is the *observability matrix* of depth *m* [27]. The rank of *P* counts the independent state directions that can be distinguished over this observation window. Its null space contains the differences between states that produce identical measurement sequences.

If rank(*P* ) = *n*, different latent states produce different delay vectors, and the inverse relation is

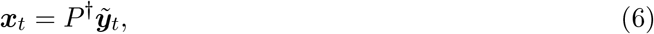

where *P* ^*†*^ is the Moore–Penrose pseudoinverse. Thus, full column rank makes the delay-coordinate map injective, providing a linear counterpart of the delay-embedding idea underlying Takens’ theorem [31]. Because *P* depends on the unknown dynamics *A*, this inverse relation is not an available reconstruction procedure. Rather, it establishes that the observation window preserves all information about the latent state. If rank(*P* ) *< n*, distinct latent states can produce the same delay vector.

Learning the dynamics also requires a trajectory that contains enough independent states to reveal their evolution. Consider a time series of latent states over *T* time steps, to which we do not have direct access:

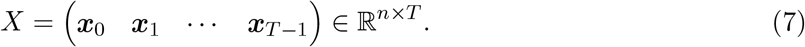

The corresponding delay vectors form the block Hankel matrix, to which we have direct access:

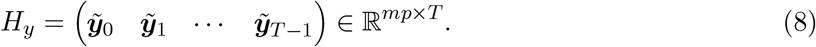

Each column is the delay vector for one starting time, with all *p* channels grouped together at each delay. This is a block Hankel matrix, whose *p ×* 1 blocks are identical along each anti-diagonal. Applying Eq. (4) to every column gives

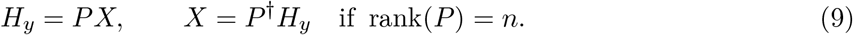

The Hankel matrix is available directly from the recordings. The inverse relation identifies when it preserves the latent trajectory information. Our objective is to recover the dynamical spectrum from this matrix without knowing *P* or the latent states themselves.

For a free response beginning at ***x***_0_, meaning evolution without further external input, the state trajectory is

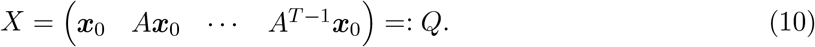

We call *Q* the *trajectory matrix*, or more specifically a Krylov trajectory matrix, because each column is obtained by applying *A* to the preceding column. The trajectory matrix *Q* also has the form of a *controllability matrix*. For a dynamical system with a control input,

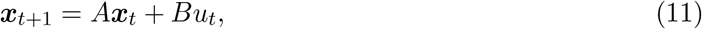

where *B* is an input matrix and *u*_*t*_ is the external input at time *t*, control theory defines the controllability matrix as

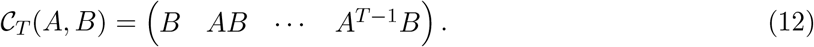

Here, if we consider a scalar impulse input of *u*_0_ = 1, *u*_*t*_ = 0 for *t >* 0, and if we set *B* = ***x***_0_ and ***x***_0_ = 0, *Q* corresponds to the controllability matrix *C*_*T*_ . The interpretation explains why *Q* can be regarded as the *controllability matrix*.

Substituting Eq. (10) into Eq. (9) yields the important factorisation of the Hankel matrix into the observability matrix *P* and the trajectory matrix *Q*,

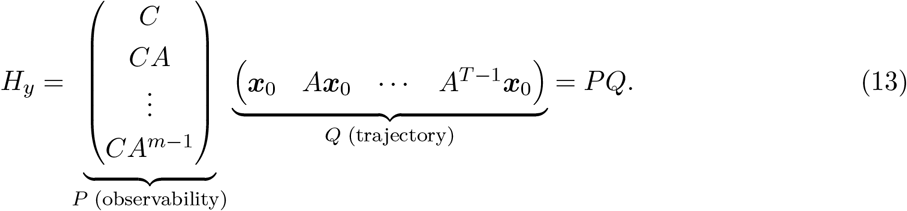

Furthermore, considering the interpretation of a controllability matrix described above, this factorisation can be interpreted as the observability–controllability factorisation of realisation theory, applied to the equivalent impulse initialisation of a free response [28, 29] (Fig. 1D).

The factorisation of the Hankel matrix clarifies two limits on extractable information from partial observation: *P* specifies which state directions the measurements can distinguish, whereas *Q* specifies which directions are present in the sampled trajectory. Consequently,

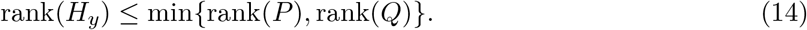

More precisely, rank(*H*_*y*_) is the dimension of the image of the sampled state subspace under *P* : it counts the independent directions in the trajectory that remain distinguishable after observation. The bound need not be an equality when both factors are rank-deficient.

The data span all *n* latent state directions in delay coordinates exactly when

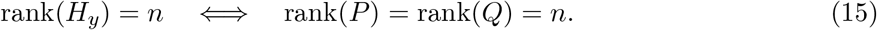

Full rank of *P* means that the measurement windows distinguish all state directions; full rank of *Q* means that the recorded trajectory spans them. This criterion refers to the true latent dimension *n*; the measured Hankel rank alone cannot exclude additional unobserved modes. We next show how Hankel DMD uses the evolution of these measurement windows to recover the dynamical spectrum directly, without first reconstructing the original neural variables.

### 2.3 Reconstructing the dynamical spectrum from Hankel DMD

Hankel DMD estimates a linear map that advances the measured delay vectors by one time step [32, 35]. Let 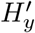 contain the same windows as *H*_*y*_, shifted forward by one sample. The paired matrices use *m* + *T* measurement samples, from time 0 through *m* + *T −* 1, and satisfy

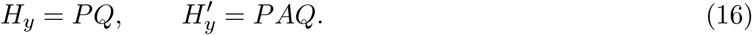

Fitting a linear map between them gives the minimum-norm least-squares estimate

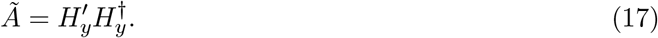

This estimate requires only the recordings; neither *P,Q*, nor *A* needs to be known.

For linear dynamics, if rank(*H*_*y*_) = *n*, equivalently rank(*P* ) = rank(*Q*) = *n*, the eigenvalues of the Hankel DMD operator *Ã* coincide with those of *A*, including their algebraic multiplicities, except for *mp − n* additional zero eigenvalues when *mp > n*. The estimated *Ã* describes the original evolution in different coordinates, which preserves its eigenvalues. If *mp > n*, the unreduced matrix *Ã* also has zero eigenvalues from the unused directions outside the data span; these are not additional latent modes. The derivation and the reduced implementation are given in Supplementary Section S1.

When the full-rank condition fails, full-system recovery is not guaranteed. Only dynamics that are both observed and excited can be recovered, and the delays and samples must suffice to distinguish them. This separates the information limit from the estimation procedure: Hankel DMD extracts a spectrum from the recording, while the observability and trajectory matrices determine whether that spectrum can represent the full system. These conditions identify the observation and trajectory limitations that the next subsection addresses.

### 2.4 Increasing recoverable information: delay embedding and perturbation

The preceding subsections establish recovery conditions for a given recording and delay depth. We now address the second question: how can changes to the observation windows and sampled trajectories increase the information available for spectral recovery?

The factorisation *H*_*y*_ = *PQ* identifies two complementary ways to improve recovery: increasing the number of state directions distinguishable through *P*, and increasing the number of independent directions represented in the trajectory matrix *Q*.

First, the rank of *P* can increase through an increase in the delay depth *m* (Fig. 1B). In neural recordings, experimental constraints often fix *C*, limiting our ability to change which variables are observed. The delay depth, however, can be increased during analysis, provided the recording contains enough samples. Additional delays append rows to *P*, allowing hidden components to become distinguishable as their effects propagate into the recorded channels. This improvement has a limit: for a linear *n*-dimensional system, the observable subspace reaches its maximum by delay depth *n*. A component that never affects the measurements remains inaccessible at every delay depth. Thus, for given dynamics, the fixed measurement matrix *C* sets a ceiling on the rank that delay embedding can achieve. For a recording of fixed length *L*, the number of paired measurement windows is *T* = *L − m*. Increasing the delay depth therefore reduces the number of columns available to sample the trajectory, so improved observability must be balanced against sufficient trajectory sampling.

Second, the rank of *Q* can increase by changing the initial state ***x***_0_ through an external perturbation. An appropriate perturbation can excite missing state directions and thereby increase the rank of *Q*. Perturbation can also re-excite rapidly decaying modes that are already present in the spontaneous trajectory in principle but have become too weak to recover reliably from finite, noisy recordings (Fig. 1C). The resulting transient response can therefore improve spectral recovery either by adding independent directions to *Q* or by increasing the amplitudes of weakly represented modes. These additional trajectory components contribute to the measured Hankel matrix only to the extent that they remain distinguishable through *P* .

This effect of perturbation on the trajectory rank can be seen directly from the state evolution. In many dissipative systems, some eigenvalues of *A* lie inside the unit circle, so their contributions decay during long periods without perturbation. Consequently, these modes can become weakly represented in the spontaneous trajectory, making *Q* poorly conditioned or effectively rank deficient in finite, noisy recordings; if a mode is absent altogether, *Q* is rank deficient even in the noise-free case. Let 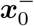 denote the initial state in the absence of perturbation, and let 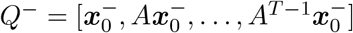 denote the corresponding spontaneous trajectory matrix. An impulse applied at time 0 changes this initial state according to

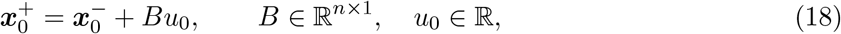

where 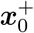 denotes the perturbed initial state, *B* specifies the state direction affected by the perturbation, and *u*_0_ its magnitude. The subsequent free response from 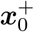 has trajectory matrix 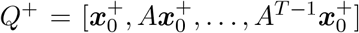. The propagation of the input direction through the dynamics is described by the finite-horizon controllability matrix

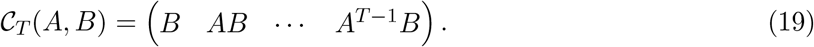

It follows directly that

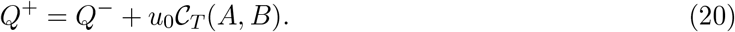

Thus, the perturbation adds the propagated input components *Bu*_0_, *ABu*_0_, …, *A*^*T −*1^*Bu*_0_ to those already present in the spontaneous trajectory. If these added components restore independent directions that are absent from *Q*^*−*^, they can increase the exact rank of *Q*^+^; if the corresponding directions are already present but only weakly represented because of decay, perturbation can instead increase their amplitudes and improve the numerical rank and conditioning of *Q*^+^ (Fig. 1C).

Exact rank is only one aspect of recovery from finite, noisy recordings. Noise can make the measured Hankel matrix full rank without adding dynamical information, so the numerical rank used in practice depends on distinguishing signal singular values from those attributable to noise. Observable and excited modes may still have very small amplitudes, or their measurement signatures may be difficult to distinguish. These limitations appear as small signal singular values and poor conditioning. Additional delays and suitable perturbations can improve practical recovery without changing the rank of the underlying noise-free Hankel matrix. Delay depth, recording length, perturbation, and retained DMD rank should therefore be evaluated together. The full-rank result establishes what is possible in principle; reliable estimation also depends on the strength and distinguishability of the measured components.

### 2.5 Extension to nonlinear dynamics through Koopman theory

Although the preceding theoretical framework assumes linear dynamics, its central arguments can also apply to nonlinear systems through Koopman operator theory. This theory describes the evolution of functions of the state using a linear operator. Importantly, the practical analysis procedure remains unchanged: the same delayed measurements are used to construct the Hankel matrices, and their evolution is estimated by the same Hankel DMD procedure. Thus, the method can be applied without first determining whether the underlying state dynamics are linear or nonlinear. What changes is the interpretation and the conditions for exact recovery. When the measured observables belong to a finite-dimensional Koopman-invariant subspace, the recovered eigenvalues can be interpreted as Koopman eigenvalues, and spectral recovery can be analysed using the same framework as in the linear case.

To make this connection precise, let *X ⊆* ℝ^*n*^ denote the state space and consider the discrete-time nonlinear system

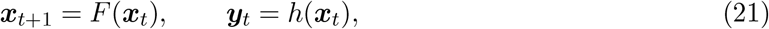

where *F* : *X → X* is the state-evolution map and *h* : *X →* ℝ^*p*^ is the measurement function. The components of *h* specify the *p* quantities measured from the state and may themselves be nonlinear. More generally, a scalar-valued function *g* : *X →* C is called an *observable*. The Koopman operator *K* acts on a function space *G* of such observables by composition with the dynamics,

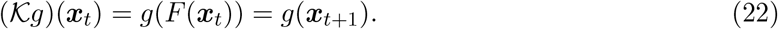

Thus, *Kg* gives the value of the observable *g* one time step after the system starts from ***x***. Although the state evolution *F* may be nonlinear, *K* is linear on *G*, satisfying *K*(*ag*_1_ + *bg*_2_) = *aKg*_1_ + *bKg*_2_ for observables *g*_1_, *g*_2_ *∈ G* and scalars *a, b*. The function space *G* is generally infinite-dimensional even when the state space is finite-dimensional [36].

Hankel DMD can recover a finite-dimensional representation of Koopman evolution when the measured observables belong to a finite-dimensional Koopman-invariant subspace and the delay coordinates and sampled trajectory sufficiently span that subspace [32]. To express this connection directly, let

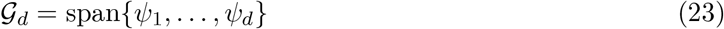

be a *d*-dimensional Koopman-invariant subspace spanned by linearly independent Koopman eigen-functions *ψ*_*i*_ : *X →* ℂ satisfying

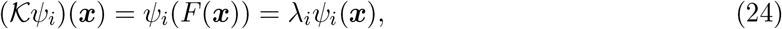

where *λ*_*i*_ *∈* ℂ is the Koopman eigenvalue associated with *ψ*_*i*_. Suppose that each component of the measurement function *h* : *X →* ℝ^*p*^ belongs to *G*_*d*_. Collecting the eigenfunctions into ***ψ*** = (*ψ*_1_, …, *ψ*_*d*_)^*⊤*^ gives

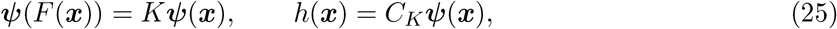

where *K* = diag(*λ*_1_, …, *λ*_*d*_) *∈* ℂ^*d×d*^ represents the Koopman evolution restricted to *G*_*d*_, and *C*_*K*_ *∈* ℂ^*p×d*^ maps the Koopman coordinates to the measured variables. Here *d* is the dimension of the invariant subspace used to represent the measurements and need not equal the latent state dimension *n*. The columns of *C*_*K*_ are the Koopman modes in the recorded channels, while ***ψ***(***x***_0_) gives their amplitudes at the initial state. The measured time series therefore evolves as

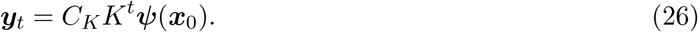

This has the same algebraic form as the linear representation ***y***_*t*_ = *CA*^*t*^***x***_0_. Consequently, the Hankel matrices factorise as

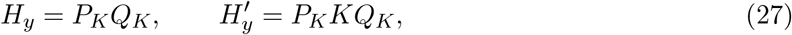

where

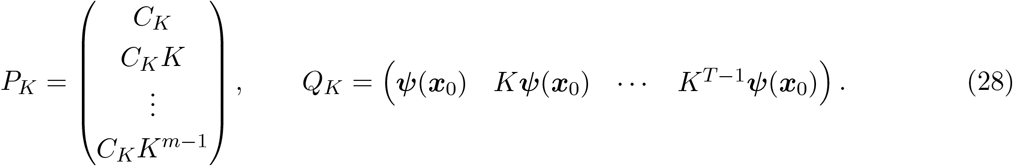

Here *P*_*K*_ *∈* ℂ^*mp×d*^ describes how the Koopman coordinates are represented across the *m* delayed measurements, whereas *Q*_*K*_ *∈* ℂ^*d×T*^ describes which directions in the invariant subspace are represented along the sampled trajectory. These factors specify the theoretical representation; neither the eigenfunctions ***ψ*** nor the matrices *K* and *C*_*K*_ need to be known beforehand. Hankel DMD instead estimates the evolution directly from the recordings as

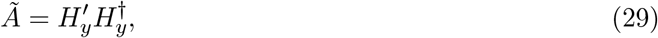

where 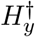 denotes the Moore–Penrose pseudoinverse. If rank(*P*_*K*_) = rank(*Q*_*K*_) = *d*, the data are noise-free, and all *d* data directions are retained, standard Hankel DMD results imply that the restriction of *A* to the data subspace has the same eigenvalues as *K* [32]. These eigenvalues, *λ*_1_, …, *λ*_*d*_, are the Koopman eigenvalues represented in this invariant subspace. Thus, as in Section 2.3, sufficient observation through *P*_*K*_ and sufficient trajectory sampling through *Q*_*K*_ allow Hankel DMD to recover the Koopman eigenvalues represented in the selected subspace. The general invariant-subspace formulation, which does not require an eigenfunction basis or a diagonalisable matrix representation, is given in Supplementary Section S1.3.

A finite-dimensional invariant subspace containing the measurements does not generally exist. Without this closure, the exact factorisation and spectral-recovery guarantee above no longer apply to a finite set of observable coordinates. Nevertheless, Hankel DMD can provide a useful finite-dimensional approximation of Koopman evolution on the sampled observables, as demonstrated in studies of delay-based representations of nonlinear dynamics [32, 37]. Accurate prediction over a finite recording window does not by itself guarantee accurate eigenvalues, and finite approximations can produce spurious modes [38]. In such approximations, the observability and trajectory matrices remain useful for understanding which components are distinguishable and sufficiently excited, although approximation error and conditioning also affect spectral accuracy. While exact recovery is harder to predict in general nonlinear systems, these principles still guide experimental design and analysis: delay embedding can improve distinguishability, and suitable perturbations can broaden the dynamics represented in the recordings.

## 3 Validation of theoretical framework

We performed both simulations and analyses of ECoG data to test the theoretical prediction that delay embedding and perturbation improve mode recovery. We first turned to a simulated linear and a nonlinear system, where the estimated eigenvalues can be compared directly with the true ones. We demonstrate that modes hidden by partial observation can be recovered by increasing the ranks of *P* and *Q* through delay embedding and perturbation, respectively. We next investigated whether the dependence on delay embedding and perturbation identified in simulation is also present in human ECoG recordings with electrical stimulation.

### 3.1 Simulations

#### 3.1.1 Both observability and trajectory determine the recovery of eigenvalues in a linear system

Raising the rank of *P* recovers eigenvalues hidden by partial observation, and raising the rank of *Q* recovers eigenvalues that were not excited in the passive trajectory. To confirm this, we considered the partially observed linear system in Eqs. (1) and (2) with ***x***_*t*_ *∈* ℝ^24^ and ***y***_*t*_ *∈* ℝ^8^, so that only 8 of the 24 state variables are observed (network and parameters in Supplementary Section S2.1, Fig. S1A, and Table S1). The system contained both slow-decaying and fast-decaying modes. The slow-decaying modes remained active along a passive trajectory, whereas the fast-decaying modes were re-excited by an impulse perturbation. We estimated the eigenvalues by DMD with no delay embedding (at delay depth of *m* = 1) and by Hankel DMD at delay depth of *m* = 8 in each case, and compared them with the true ones. Increasing the delay depth raised the rank of *P* from 8 to 24 (Fig. 2A), whereas the impulse perturbation raised the rank of *Q* from 10 to 24 (Fig. 2B).

**Figure 2.**
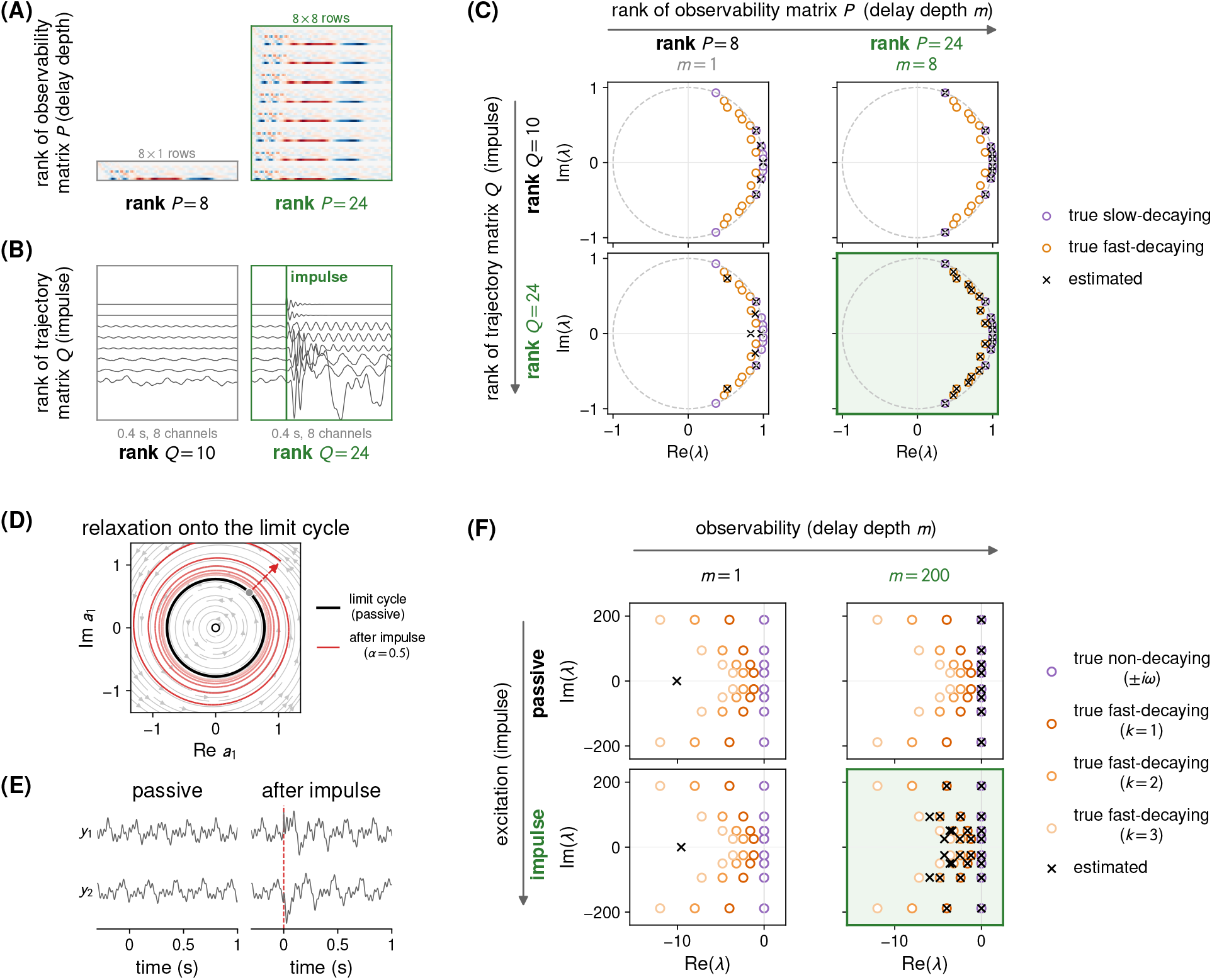
Delay embedding and perturbation relieve complementary rank bottlenecks. **(A)** Increasing the delay depth increases the observability rank *P* . **(B)** Impulse perturbation increases the excitation rank *Q*. **(C)** Eigenvalue recovery in the discrete-time complex plane for combinations of the rank of the observability matrix *P* and the rank of the trajectory matrix *Q*. The dashed curve indicates the unit circle. Purple circles denote true slow-decaying modes, orange circles denote true fast-decaying modes, and crosses denote estimated eigenvalues. **(D–F)** The same two levers on a nonlinear system: four uncoupled Stuart–Landau oscillators (4, 8, 15 and 30 Hz; Eq. (30)), observed through two linear projections of their eight-dimensional state. **(D)** Flow of the 4 Hz oscillator (grey streamlines). The passive trajectory runs on the limit cycle (black); an impulse (dashed arrow, from the grey dot) displaces the state, which then spirals back onto the cycle (red, fading with time). For visibility this panel uses *α* = 0.5; the spectra in (F) use *α* = 0.2. The open circle marks the unstable fixed point at the origin. **(E)** The two observed projections *y*_1_ and *y*_2_ of the passive trajectory (left) and of the perturbed trajectory (right); the dashed line marks the impulse at *t* = 0, after which the amplitude swells and relaxes back within about 0.5 s. **(F)** DMD spectra with delay depth (observability) on the columns and the impulse (*α* = 0.2; excitation) on the rows. Circles are the true non-decaying (*± iω*) and fast-decaying ( *−*2*kr ± iω, k* = 1, 2, 3) eigenvalues, and crosses are the DMD estimates (real part in s^*−*1^, imaginary part in rad s^*−*1^).

Without delay embedding and without perturbation, DMD recovered only five eigenvalues close to the true slow-decaying modes (Fig. 2C, upper left). Purple and orange open circles indicate the true eigenvalues of the slow-decaying and fast-decaying modes, respectively. Black crosses indicate the DMD estimates. The eight observed variables were not sufficient to recover the true modes. This failure occurred because the fast-decaying modes were not excited in the passive trajectory and the oscillations of the unobserved state variables could not be reconstructed from the partially observed signals.

Delay embedding enabled recovery of all 10 slow-decaying eigenvalues from the passive trajectory, while the fast-decaying modes remained unrecovered (Fig. 2C, upper right). Increasing the delay depth from *m* = 1 to *m* = 8 raised the rank of *P* from 8 to the full latent dimension of 24, so that the latent state could be reconstructed uniquely from the delay vector. The rank of *Q* remained 10, because no external stimulus was applied and the fast-decaying modes were still not excited. Delay embedding enabled recovery of the slow-decaying modes, but could not recover fast-decaying modes.

Impulse perturbation enabled recovery of five fast-decaying eigenvalues (Fig. 2C, lower left). The impulse perturbation re-excited the fast-decaying modes, raising the rank of *Q* from 10 to the full latent dimension of 24. In addition to recovering three slow-decaying eigenvalues, the perturbation re-excited the fast-decaying modes and enabled recovery of the fast-decaying eigenvalues. If we do not use delay embedding, the rank of *P* remained 8 despite the perturbation. Impulse perturbation supplied the excitation that the passive trajectory lacked, but could not compensate for the limited observability matrix rank.

With both delay embedding and impulse perturbation, Hankel DMD recovered all 24 eigenvalues (Fig. 2C, lower right). The perturbation and delay embedding raised the ranks of *Q* and *P* to 24, respectively. This means that the trajectory contained every mode and the delay vectors distinguished every latent state direction. Full recovery was achieved only when both conditions were satisfied. These results demonstrate the complementary roles of observation and excitation, consistent with the full-rank recovery condition in Eq. (15) derived in Section 2.2.

The results above were obtained in a representative setting designed to expose the effects of observability and excitation as clear rank bottlenecks. We therefore examined how eigenvalue recovery changed when these idealized conditions were relaxed by varying the delay depth and perturbation site, and by introducing recurrent connections, noise, and different perturbation amplitudes (Supplementary Sections S2.1.2 and S2.1.3). Across these settings, recovery continued to depend on both the information available through observation and the extent to which the underlying dynamics were excited.

#### 3.1.2 Extension to a nonlinear system

We next investigated whether the dependence on observation and excitation identified in the linear system also extends to nonlinear dynamics. We considered four uncoupled Stuart–Landau oscillators,

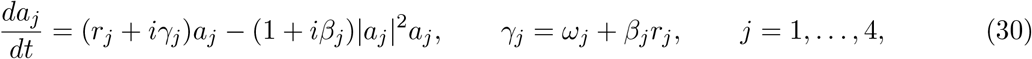

where *a*_*j*_ *∈* ℂ is the complex amplitude of oscillator *j* (Fig. 2D). For *r*_*j*_ *>* 0, each oscillator approaches a stable limit cycle with radius 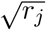 and angular frequency *ω*_*j*_. When displaced from the cycle, it exhibits transient amplitude dynamics and gradually returns to the cycle.

We focused on two families of Koopman eigenvalues corresponding to the slow- and fast-decaying eigenvalues of the linear simulation:

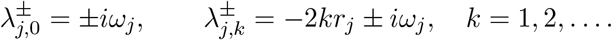

A continuous-time Koopman eigenvalue *λ* describes a contribution evolving as *e*^*λt*^: its real part determines growth or decay, and its imaginary part determines oscillation frequency. Thus, the eigenvalues 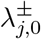 describe non-decaying oscillations along the cycle and correspond to the slow-decaying eigenvalues of the linear system. These eigenvalues are expressed in the passive trajectory without perturbation. The second family 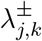 describes progressively faster-decaying contributions to the relaxation after displacement, and we refer to these as fast-decaying eigenvalues. The sequence arises because products of Koopman eigenfunctions add their continuous-time eigenvalues: multiplying the *k*th power of a radial eigenfunction with eigenvalue *−*2*r*_*j*_ by a phase eigenfunction with eigenvalue *±iω*_*j*_ gives *−*2*kr*_*j*_ *± iω*_*j*_.

For trajectories sufficiently close to the limit cycle, the oscillator state can be expressed as

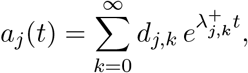

where the coefficients *d*_*j,k*_ depend on the initial state (derivation in Supplementary Section S2.2). On the cycle, all coefficients with *k ≥* 1 vanish, leaving only the non-decaying oscillation. Away from the cycle, infinitely many terms contribute, so any finite-dimensional representation is approximate. Near the cycle, higher-order contributions become small, and a finite set of eigenvalues can approximate the measured dynamics. Rather than attempting to recover the full infinite spectrum, we tested whether the non-decaying eigenvalues and the first three orders of fast-decaying eigenvalues (*k* = 1, 2, 3) became recoverable as observation and excitation were increased.

As in the linear simulation, we independently varied perturbation and delay embedding under partial observation. The real and imaginary parts of the four oscillator states formed an eight-dimensional state ***x***, observed through two fixed random linear mixtures, ***y*** = *C****x***, with *C ∈* ℝ^2*×*8^ (Fig. 2E). In the passive condition, the trajectory remained on the limit cycles, so the fast-decaying modes were not excited. In the perturbed condition, an impulse displaced the state according to ***x***(0^+^) = ***x***(0^*−*^) + *α****b***, with a fixed direction ***b*** and *α* = 0.2, exciting transient relaxation towards the cycles. In each condition, we estimated eigenvalues using standard DMD without delay embedding (*m* = 1) and Hankel DMD with delay embedding (*m* = 200).

Delay embedding was required to recover eigenvalues from the two-channel observations (Fig. 2F). Without delay embedding, DMD recovered none of the target eigenvalues in either the passive or perturbed condition (Fig. 2F, left column): the two instantaneous measurements were insufficient to separate the dynamics of the four oscillators. Delay embedding recovered the non-decaying eigenvalue pair *±iω*_*j*_ for every oscillator from the passive trajectory (Fig. 2F, upper right). However, it did not recover the fast-decaying eigenvalues because these modes were not excited while the trajectory remained on the cycles.

Only when perturbation was combined with delay embedding were the fast-decaying eigenvalues recovered (Fig. 2F, lower right). The *k* = 1 pair was recovered for all four oscillators and the *k* = 2 pair for three of the four, while the *k* = 3 pair was not recovered; the non-decaying eigenvalues remained recovered for all four oscillators (Fig. 2F, lower right). Recovery of fast-decaying eigenvalues after perturbation was also observed across other perturbation magnitudes and observation matrices at *m* = 200 (Supplementary Table S2B). The nonlinear system therefore showed the same complementary dependence as the linear system. Perturbation supplied the missing excitation, whereas delay embedding supplied the temporal information needed to resolve the excited dynamics from partial observations.

### 3.2 Analysis of human ECoG data

We next investigated whether delay embedding and perturbation, the manipulations that raise the ranks of the observability and trajectory matrices in theory, changed the validated eigenvalue estimates in human ECoG recordings as predicted. We analysed a publicly available ECoG dataset acquired during repeated single-pulse electrical stimulation [34]. The recordings were well suited for assessing the effects of perturbation because each trial contained both pre-stimulus spontaneous activity and post-stimulus perturbed activity from the same channels. In addition to comparing conditions with and without perturbation, the depth of the delay embedding can also be varied to examine the effect of observability.

Because the true eigenvalues are unknown in human data, mode recovery could not be evaluated directly against ground truth as in the simulations. Instead, we examined how perturbation and delay embedding changed the number of the estimated eigenvalues by comparing pre- and post-stimulus activity and varying the delay depth, respectively. Importantly, to ensure the reliability of the estimated eigenvalues, we checked whether each eigenpair estimated from one trial was supported by other trials from the same stimulation site (see Methods and Supplementary Section S3.2). We refer to eigenpairs that passed this validation criterion as validated estimates. Then, we evaluated whether the number of only such “validated” estimates of eigenvalues increased. We first characterised these effects in one participant at one stimulation site and then examined their consistency across participants.

#### 3.2.1 Perturbation and delay embedding reveal higher-frequency and fast-decaying eigenvalue estimates in an example participant

At a single stimulation site in an example participant, delay embedding and perturbation changed different properties of the validated eigenvalue estimates. Delay embedding increased the number of validated eigenvalue estimates and extended them to higher oscillation frequencies, whereas perturbation increased the number of validated estimates of fast-decaying eigenvalues. The ECoG electrodes covered the cortical surface, and stimulation was delivered through a pair of adjacent electrodes (Fig. 3A). We analysed the pre-stimulus window (*−*310 to *−*10 ms) and the post-stimulus window (10 to 310 ms) around each stimulation (Fig. 3B). For each window and each delay depth, we fitted DMD to each of the ten stimulation trials separately, tested every resulting eigenpair on the other nine trials (Supplementary Fig. S3A–D shows the candidate eigenpairs removed by each criterion at this site), and pooled the validated eigenvalue estimates across the ten fits. Because the same eigenvalue can be recovered from several trials, these counts are numbers of validated estimates, not numbers of distinct eigenvalues. We classified eigenvalues with decay rates at or below *−*10 s^*−*1^, corresponding to time constants of 100 ms or less, as fast-decaying (orange), and the remaining eigenvalues as slow-decaying (purple). This threshold provides a descriptive split of a continuous distribution (Supplementary Fig. S4).

**Figure 3.**
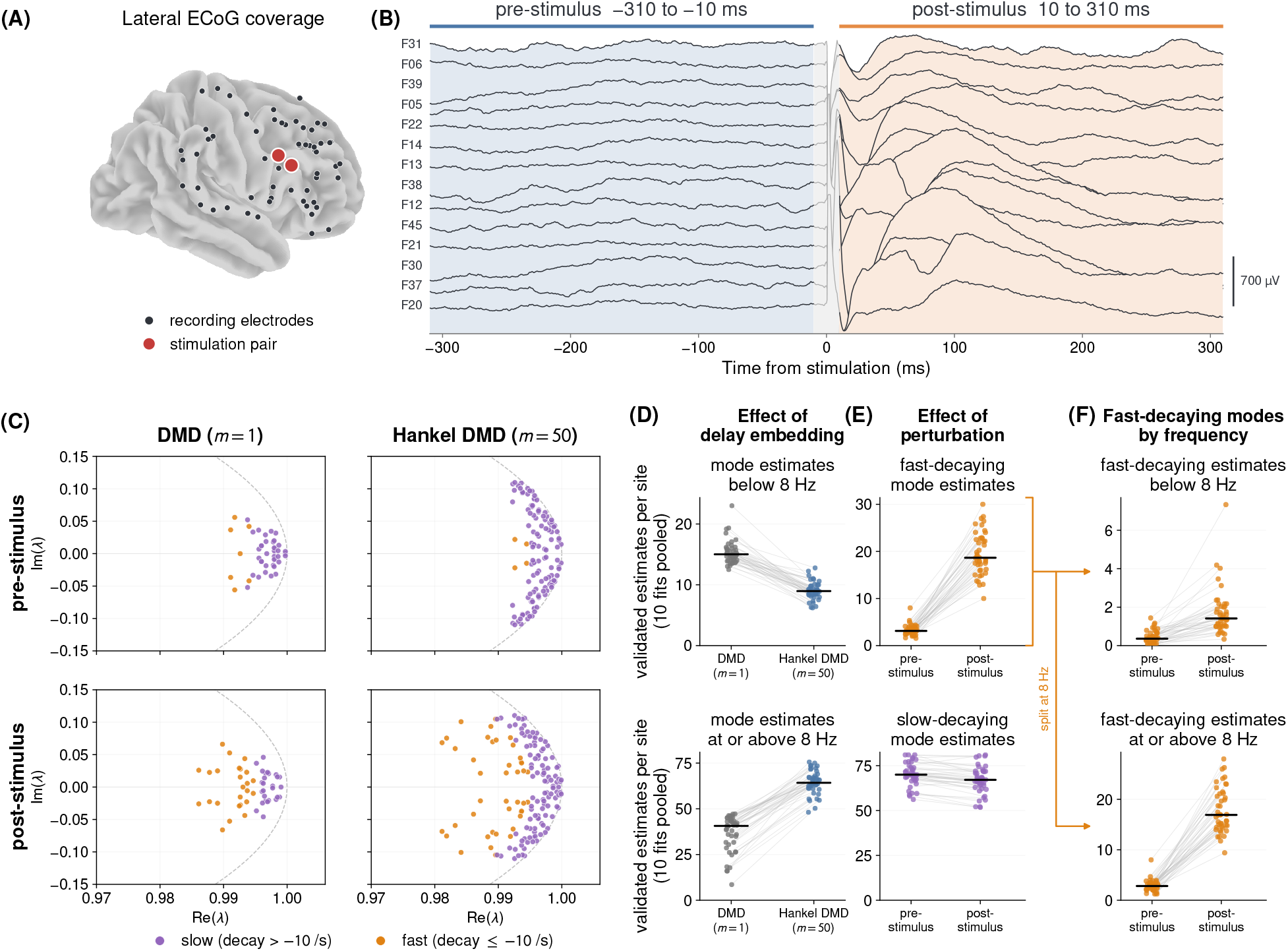
Perturbation adds validated estimates of fast-decaying eigenvalues, whereas delay embedding adds higher-frequency estimates in human ECoG. **(A)** Lateral ECoG coverage in an example participant. Black circles indicate recording electrodes, and red circles indicate the stimulation pair. **(B)** Example ECoG signals around electrical stimulation. Blue and orange regions indicate the pre-stimulus ( *−*310 to *−*10 ms) and post-stimulus (10 to 310 ms) windows used for analysis, respectively. **(C)** Validated eigenvalue estimates from the example participant at a single stimulation site in the discrete-time complex plane. Columns compare standard DMD without delay embedding (*m* = 1) and Hankel DMD with delay embedding (*m* = 50). Rows compare pre- and post-stimulus windows. The dashed curve indicates the unit circle. Orange points denote estimates of fast-decaying eigenvalues (decay *≤−*10 s^*−*1^), whereas purple points denote estimates of slow-decaying eigenvalues (decay *> −*10 s^*−*1^). **(D–F)** Across-participant comparison of validated eigenvalue-estimate counts. Each point represents one participant (*n* = 44), and the count is the number of validated estimates at a site, pooled over its ten per-trial fits as in (C), averaged over that participant’s stimulation sites. Grey lines connect paired estimates from the same participant, and black horizontal bars indicate the median. **(D)** Numbers of validated estimates in the pre-stimulus window below 8 Hz (top) and at or above 8 Hz (bottom), comparing DMD (*m* = 1) and Hankel DMD (*m* = 50). **(E)** Numbers of validated estimates of fast-decaying (top) and slow-decaying (bottom) eigenvalues, comparing pre- and post-stimulus windows using Hankel DMD. **(F)** Numbers of validated estimates of fast-decaying eigenvalues below 8 Hz (top) and at or above 8 Hz (bottom), comparing pre- and post-stimulus windows using Hankel DMD. The orange guide lines indicate that (F) splits the fast-decaying estimates of (E, top) at 8 Hz.

First, we showed that delay embedding alone increased the total number of validated estimates of slow-decaying eigenvalues and extended them to higher oscillation frequencies (Fig. 3C, upper row). In the pre-stimulus window, increasing the delay depth from *m* = 1 to *m* = 50 raised the number of validated eigenvalue estimates from 24 to 63. Almost all of the added estimates were of slow-decaying eigenvalues, whose number increased from 20 to 61. The added estimates spread along the unit circle to larger angles, that is, to higher oscillation frequencies. Without perturbation, delay embedding therefore yielded more validated eigenvalue estimates over a broader frequency range.

Next, we confirmed that perturbation increased the number of validated estimates of fast-decaying eigenvalues (Fig. 3C, left column). With standard DMD (*m* = 1), the number of validated estimates of fast-decaying eigenvalues increased from four of 24 in the pre-stimulus window to 13 of 28 in the post-stimulus window, whereas the number of validated estimates of slow-decaying eigenvalues changed little, from 20 to 15. The added estimates appeared as orange points far inside the unit circle. Without delay embedding, perturbation therefore changed the decay composition of the validated eigenvalue estimates, while their frequency range remained similar.

Combining delay embedding and perturbation increased the number of validated estimates of both slow-decaying and fast-decaying eigenvalues (Fig. 3C, lower right). Hankel DMD (*m* = 50) applied to the post-stimulus window yielded 79 validated eigenvalue estimates, of which 57 were of slow-decaying eigenvalues and 22 were of fast-decaying eigenvalues. Compared with delay embedding alone (upper right), the number of validated estimates of slow-decaying eigenvalues was maintained, from 61 to 57, and that of fast-decaying eigenvalues increased from 2 to 22. Compared with perturbation alone (lower left), the number of validated estimates of slow-decaying eigenvalues increased from 15 to 57, and that of fast-decaying eigenvalues increased from 13 to 22. The fast-decaying eigenvalues also spread to larger angles than at *m* = 1, extending to higher oscillation frequencies. Delay embedding and perturbation thus played complementary roles, as in the simulations (Fig. 2), and their combination yielded many validated estimates of both slow-decaying and fast-decaying eigenvalues.

#### 3.2.2 Consistency of perturbation and delay embedding effects across participants

Across 44 participants and 1318 stimulation sites (see Supplementary Section S3.1 for data selection), delay embedding extended the range of validated eigenvalue estimates to higher frequencies and perturbation increased the number of validated estimates of fast-decaying eigenvalues. To compare participants, we averaged the number of validated estimates per site over each participant’s stimulation sites, so each point in Fig. 3D–F represents one participant.

Delay embedding yielded additional validated eigenvalue estimates at higher frequencies (Fig. 3D). We first compared the validated eigenvalue estimates obtained from pre-stimulus (passive) activity with standard DMD (*m* = 1) and Hankel DMD (*m* = 50). Hankel DMD yielded fewer validated eigenvalue estimates below 8 Hz than standard DMD, but many more at or above 8 Hz. This pattern matches the upper row of Fig. 3C, where delay embedding extended the validated eigenvalue estimates to larger angles in the complex plane.

Perturbation increased the number of validated estimates of fast-decaying eigenvalues while leaving that of slow-decaying eigenvalues nearly unchanged (Fig. 3E). To assess perturbation under delay embedding, as in the right column of Fig. 3C, we fixed the delay depth at *m* = 50 and compared the pre- and post-stimulus windows. The number of validated estimates of fast-decaying eigenvalues (decay rate *≤ −*10 s^*−*1^) increased after stimulation in all 44 participants, from a median of 3.1 to 18.7 per site, with a median within-participant increase of 15.5 (Fig. 3E, top). The number of validated estimates of slow-decaying eigenvalues (decay rate *> −*10 s^*−*1^) barely changed, from a median of 70.0 to 67.1, with a median within-participant change of *−*3.5 (Fig. 3E, bottom). The increase after stimulation was therefore confined to fast-decaying eigenvalues. This result was robust to the rank and the validation criteria (Supplementary Section S3.4 and Fig. S5).

The increase in validated estimates of fast-decaying eigenvalues after stimulation lay predominantly at higher frequencies (Fig. 3F). We divided the validated estimates of fast-decaying eigenvalues in Fig. 3E (top) into those below 8 Hz and those at or above 8 Hz (continuous distribution in Supplementary Fig. S4). Their number increased after stimulation in all 44 participants in both ranges, but the increase was much larger at or above 8 Hz, with a median within-participant increase of 13.9 per site compared with 1.0 below 8 Hz.

These results are consistent with the distinct effects of delay embedding and perturbation predicted by the theoretical framework. Delay embedding enriches the observability matrix, allowing higher-frequency eigenvalues to be resolved from the same pre-stimulus activity. Perturbation enriches the trajectory matrix by driving the system along fast-decaying directions that are weakly excited before stimulation, consistent with perturbation revealing transient dynamics that were weakly represented before stimulation. Together with the simulations, these findings show that delay embedding and active perturbation reveal components of latent neural dynamics that are poorly captured by passive recordings.

## 4 Discussion

Understanding how interacting neural populations communicate requires inferring their dynamics from partial observations. We used control theory to show that mode recovery depends on both observation and excitation. Writing the Hankel matrix as *H*_*y*_ = *PQ* is the observability–controllability factorisation of realisation theory [27–30]. The rank bound is what makes the two constraints complementary rather than interchangeable. Delay embedding and perturbation address these two constraints, respectively. In the linear simulation, combining the two recovered all eigenvalues, whereas neither alone did. In human ECoG recordings, stimulation increased the number of validated estimates of fast-decaying eigenvalues, while delay embedding yielded additional validated estimates of eigenvalues at higher frequencies. These results show why both observation and excitation need to be considered when studying neural dynamics from partial recordings.

The physiological role of the additional validated estimates of eigenvalues that our analysis revealed remains open. Our analysis shows that perturbation and delay embedding change which estimates pass cross-trial validation, but not which neural processes these estimates represent. Fast-decaying estimates revealed by perturbation may reflect components that are weakly expressed during spontaneous activity, whereas higher-frequency estimates revealed by delay embedding may reflect dynamics that cannot be separated from instantaneous observations. Whether these estimates of eigenvalues correspond to different neural pathways, spatial patterns, or functional processes remains to be tested. Future work should examine the spatial structure and physiological properties of the validated estimates of eigenvalues and determine what information perturbation and delay embedding add to the analysis of neural dynamics.

The framework also gives a principle for experimental design. Stimulation sites determine which modes are excited, and recording sites determine which of the excited modes are observed. Since recovery is limited by whichever factor is weaker, the two should be chosen together. Previous studies have designed perturbations to improve system identification [26] and used adaptive stimulation to infer connectivity [33]. Our framework adds recording sites as a second design variable. Interventional experiments that vary stimulation and recording sites together could test whether the validated estimates change as the framework predicts.

## 5 Methods

### 5.1 Linear simulation

We tested the theory of Section 2 on a linear system whose eigenvalues of *A* are known exactly. Writing 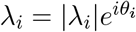, the magnitude and phase of each eigenvalue characterise the temporal behaviour of the corresponding mode. Modes with |*λ*_*i*_| *<* 1 decay, those with |*λ*_*i*_| = 1 neither grow nor decay, and those with |*λ*_*i*_| *>* 1 are unstable. In the Re(*λ*)–Im(*λ*) plane, these modes therefore lie inside, on, and outside the unit circle, respectively. The phase *θ*_*i*_ gives the phase advance per sample and corresponds to an oscillation frequency *f*_*i*_ = |*θ*_*i*_|*/*(2*π*Δ*t*) for a sampling interval Δ*t*.

We used this representation to construct a system with *n* = 24 state variables forming 12 damped oscillators, sampled at 500 Hz. Oscillator *k* was represented by a 2 *×* 2 block *ρ*_*k*_*R*(*θ*_*k*_), whose eigenvalues are 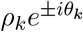. Thus, *ρ*_*k*_ directly set the decay rate of the corresponding oscillatory mode, whereas *θ*_*k*_ = 2*πf*_*k*_*/*500 set its oscillation frequency. We assigned *ρ*_*k*_ = 0.999 to five oscillators to produce slow-decaying modes and *ρ*_*k*_ *∈* [0.89, 0.96] to seven oscillators to produce fast-decaying modes. Frequencies spanned 4–95 Hz and were assigned independently of *ρ*_*k*_, so that the angular position of an eigenvalue did not reveal whether its magnitude corresponded to a slow-decaying or fast-decaying mode. The oscillators were coupled by 23 directed 2 *×* 2 blocks (gains 0.12–0.36) forming a directed acyclic graph. Every edge pointed from a lower- to a higher-indexed oscillator, making *A* block lower-triangular; consequently, coupling altered the interactions among oscillators without changing the eigenvalues set by the diagonal blocks. We observed *p* = 8 of the 24 state variables. Full specification is in Supplementary Section S2.1.

We independently varied perturbation and delay embedding, yielding four conditions. Perturbation was either absent or applied as a single impulse, and delay embedding was performed with either *m* = 1 or *m* = 8. Each run started with a 200-sample (0.4 s) settling segment, and the following 350 samples (0.7 s) were analysed. The main simulation contained no noise. For the perturbation condition, an impulse of amplitude *α* = 3 was added to the first coordinate of one oscillator at the end of the settling segment. Eigenvalues were estimated using DMD or Hankel DMD (Eq. (17)) with the truncation of Supplementary Eq. (S7), keeping the singular values of *H*_*y*_ above 10^*−*10^*σ*_1_, where *σ*_1_ is the largest singular value, so that the truncation rank equalled the numerical rank of *H*_*y*_. Recovery was then assessed by optimally matching the estimated and true eigenvalues using the Hungarian algorithm. A true eigenvalue was counted as recovered when its assigned estimate lay within 10^*−*2^.

### 5.2 Nonlinear simulation

We simulated the four uncoupled Stuart–Landau oscillators of Eq. (30) (Section 3.1.2). The parameter *β*_*j*_ makes the rotation speed depend on the amplitude, and *γ*_*j*_ = *ω*_*j*_ + *β*_*j*_*r*_*j*_ was chosen so that the angular frequency on the limit cycle equals *ω*_*j*_. Parameter values are listed in Supplementary Table S2A.

The observations are real linear combinations of Re *a*_*j*_ and Im *a*_*j*_. Following the Koopman expansion of a Stuart–Landau oscillator [39], they therefore contain only the eigenvalues 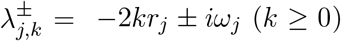 defined in Section 3.1.2. Harmonics of *ω*_*j*_ are absent because the dynamics are invariant under rotation of each *a*_*j*_, and combinations across oscillators are absent because the oscillators are uncoupled (Supplementary Section S2.2). The expansion holds whenever |*a*_*j*_|^2^ *> r*_*j*_*/*2. As the target set for recovery, we used the non-decaying eigenvalues (*k* = 0) and the fast-decaying eigenvalues of orders *k* = 1, 2, 3 for all four oscillators, 32 eigenvalues in total.

The real and imaginary parts of the four amplitudes formed the state ***x*** *∈* ℝ^8^, which was observed as ***y*** = *C****x*** through a fixed random matrix *C ∈* ℝ^2*×*8^ with unit-norm rows. In the passive condition, each oscillator started on its limit cycle at a random phase. In the perturbed condition, the impulse ***x***(0^+^) = ***x***(0^*−*^) + *α****b*** was applied at *t* = 0, with ***b*** the all-ones vector and *α* = 0.2. In each condition, eigenvalues were estimated from the same observed time series by standard DMD (*m* = 1) and Hankel DMD (*m* = 200), converted to continuous time, and matched one-to-one to the target eigenvalues. Sampling, rank selection, filtering, and the recovery criteria are given in Supplementary Section S2.2.

### 5.3 ECoG dataset and preprocessing

We analysed the public dataset ds004080, in which subdural ECoG was recorded during clinical single-pulse electrical stimulation [34]. It contains 74 participants aged 4–51 years, each with 40–111 good recording channels. Pulses were delivered to pairs of adjacent contacts while the participant rested.

We retained runs sampled at 2048 Hz with monophasic, 8 mA, 1 ms pulses, and included only pulses separated from both the preceding and following pulses by at least 4.75 s to minimise overlap between responses. A participant–site pair became a candidate when one run contained at least ten such pulses. Channels and trials with amplifier plateaus, persistent offsets, or flat or non-finite samples were excluded, and ten passing trials, evenly spaced in time, were taken from each pair (Supplementary Section S3.1 and Table S3). These quality-control criteria yielded 1503 stimulation sites from 50 participants. Requiring every fit at a site, for both methods and both analysis windows, to support the fixed rank *r* = 50, as described below, left 1318 sites from 44 participants for the primary analysis.

The only preprocessing was removal of each channel’s mean within each window, with no resampling, filtering, or re-referencing. The pre-stimulus window spanned *−*310 to *−*10 ms and the post-stimulus window +10 to +310 ms, identical in length. The 10 ms gap on each side of the pulse excluded the stimulation artefact.

### 5.4 Hankel DMD estimation

Each fit used a single trial and a single window. For *p* channels and *L* samples, we stacked *m* consecutive samples into each column of a block Hankel matrix with *T* + 1 columns, *T* = *L − m*. Dropping its last or its first column gave the Hankel matrices *H*_*y*_ and 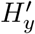 of Section 2.3, each in R^*mp×T*^, which we write as the snapshot pair 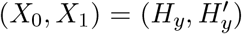.The depth *m* = 1 is standard DMD on the raw channels. The analysis compared standard DMD (*m* = 1) with Hankel DMD (*m* = 50).

We projected the snapshots onto the leading *r* left singular vectors *Û* of *X*_0_, giving *Z*_0_ = *Û*^*\**^*X*_0_ and *Z*_1_ = *Û*^*\**^*X*_1_. The reduced Hankel DMD operator *Ã*_*r*_ was the least-squares solution of *Z*_1_ *≈ Ã*_*r*_*Z*_0_ [10]. Each candidate eigenpair consists of an eigenvalue *λ*_*j*_ of *Ã*_*r*_ and its left eigenvector ***w***_*j*_, with 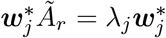. We use left rather than right eigenvectors because the row vector 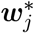 *Z*_0_ gives the values of the corresponding eigenfunction at the sampled times, which can be evaluated on held-out trials (Section 5.5). The continuous-time eigenvalue *ν*_*j*_ = log *λ*_*j*_*/*Δ*t*, with Δ*t* = 1*/*2048 s, gives the frequency *f*_*j*_ = Im *ν*_*j*_*/*(2*π*) and the decay rate Re *ν*_*j*_ = log |*λ*_*j*_|*/*Δ*t*. We kept one member of each conjugate pair (*f*_*j*_ *≥* 0).

The rank was fixed at *r* = 50 rather than selected per window. Because standard DMD has rank at most *p*, this requirement excluded sites with fewer than 50 good channels. Refitting at rank 30, which retains all 1503 quality-controlled sites from 50 participants, gave the same result (Supplementary Section S3.4). A data-driven rank grows with signal-to-noise ratio, which differs between the pre- and post-stimulus windows, and would confound any comparison of the number of validated estimates of eigenvalues. The rank fell below 50 only when fewer singular values exceeded 10^*−*4^ *σ*_1_ or when *mp <* 50. Sites where any fit fell below 50 were excluded from the primary comparison (Supplementary Section S3.1).

### 5.5 Cross-trial validation of candidate eigenpairs

A candidate eigenpair counts as validated only if it also describes trials it was not fitted on. For each participant, site, window, and method, we fitted *Û, λ*_*j*_, and ***w***_*j*_ on one of the ten trials and froze them. Nothing was re-estimated on the other nine. For held-out trial *q* we projected that trial’s own snapshots onto the frozen basis, giving 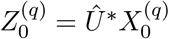 and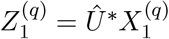, and computed the eigenfunction values 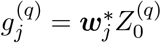 and 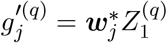. The residual measures how far the eigenvalue relation fails on that trial [38, 40],

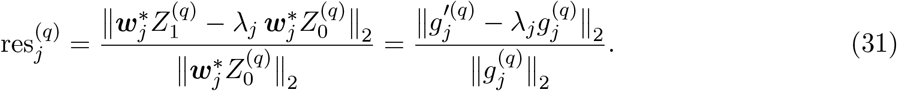

The residual in Eq. (31) checks one time step and is therefore insensitive to errors in |*λ*_*j*_| that accumulate over many steps. A second check, multi-step decay validation, compares the decay over a lag of *ℓ* = 10 samples (4.9 ms) with the decay the data show. On each held-out trial we computed the *ℓ*-step decay factor of the eigenfunction,

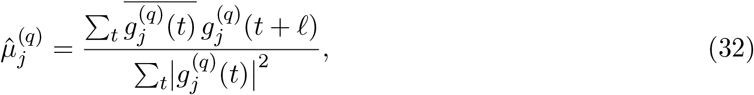

and excluded a candidate eigenpair when 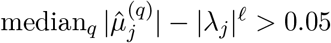, that is, when its fitted decay was clearly faster than the decay of its eigenfunction on held-out data (selection at an example stimulation site and synthetic calibration in Supplementary Fig. S3).

A candidate eigenpair was validated when it met four conditions. Its frequency was at most 100 Hz (0 *≤ f*_*j*_ *≤* 100 Hz), it was stable (|*λ*_*j*_| *<* 1), its held-out residual was at most 0.1 in at least five of the nine held-out trials (equivalently, a median residual of at most 0.1), and it passed multi-step decay validation. Validated estimates were grouped as fast-decaying (decay rate at or below *−*10 s^*−*1^) or slow-decaying, and as low-frequency (below 8 Hz) or high-frequency (at or above 8 Hz). The joint frequency–decay distribution without these thresholds is shown in Supplementary Section S3.3 and Supplementary Fig. S4.

### 5.6 Across-participant statistics

Each participant carries equal weight. For one site, one window, and one method, we pooled the validated estimates of the ten single-trial fits, as in Fig. 3C, giving the number of estimates per site. We then averaged this quantity over a participant’s stimulation sites. Contrasts are paired within participant, because the pre- and post-stimulus windows come from the same trials of the same site. We report the median across participants of these values and of the paired within-participant differences, together with the number of participants changing in each direction. The supplement gives means over participants, each weighted equally, with descriptive 95% bootstrap intervals over participants. We ran no hypothesis tests and describe the distribution across participants instead (Supplementary Section S3.4 and Supplementary Fig. S5).

## Data and code availability

The intracranial recordings analysed here are public under CC0 as OpenNeuro dataset ds004080, https://openneuro.org/datasets/ds004080 [34]. All analysis and simulation code are publicly available at https://github.com/RakuMiyoshi/dmd-partial-observation under the MIT licence, together with the per-site configuration manifests and the summary tables behind every figure.

## Acknowledgements

We thank members of the laboratory for repeated critical discussions of the manuscript, and the contributors to OpenNeuro dataset ds004080 for making the recordings public. D.S. is supported by RIKEN Center for Brain Science, the Special Postdoctoral Research Program at RIKEN. This work was supported by JSPS KAKENHI Grant Number JP23H04834 and JST Moonshot R&D Grant Number JPMJMS2012.

