## Supplementary Material for "Recovering Hidden Neural Oscillatory Modes from Partial Observations through Delay Embedding and Perturbation"

### S1 Mathematical details of state reconstruction and Hankel DMD

This section provides the mathematical details for the state-reconstruction and dynamical-spectrum results in the main text. We consider the linear dynamics and observations $\boldsymbol{x}_{t+1} = A\boldsymbol{x}_t$ and $\boldsymbol{y}_t = C\boldsymbol{x}_t$ in Eqs. (1) and (2), where $\boldsymbol{x}_t \in \mathbb{R}^n$ is the latent state, $A \in \mathbb{R}^{n\times n}$ is the state-evolution matrix, $\boldsymbol{y}_t \in \mathbb{R}^p$ is the recorded activity, and $C \in \mathbb{R}^{p\times n}$ is the observation matrix, with $p \leq n$.

The Hankel factorisation separates the two requirements for the delay data to represent all latent state directions. For delay depth $m$ and $T$ snapshot pairs, define the observability matrix $P \in \mathbb{R}^{mp\times n}$ and trajectory matrix $Q \in \mathbb{R}^{n\times T}$ by

$$P = \begin{pmatrix} C \\ CA \\ \vdots \\ CA^{m-1} \end{pmatrix}, \qquad Q = \begin{pmatrix} \boldsymbol{x}_0 & A\boldsymbol{x}_0 & \cdots & A^{T-1}\boldsymbol{x}_0 \end{pmatrix}. \tag{S1}$$

These are the matrices introduced in Eqs. (5) and (10) of the main text. The Hankel matrices $H_y, H_y' \in \mathbb{R}^{mp\times T}$ satisfy $H_y = PQ$ and $H_y' = PAQ$, as in Eqs. (13) and (16), and use $m+T$ measurement samples from time 0 through $m+T-1$. Because $P$ and $Q$ each have rank at most $n$, $\operatorname{rank}(H_y) = n$ requires $\operatorname{rank}(P) = \operatorname{rank}(Q) = n$. Conversely, if $P$ has full column rank, multiplication by $P$ preserves the rank of $Q$, so $\operatorname{rank}(P) = \operatorname{rank}(Q) = n$ implies $\operatorname{rank}(H_y) = n$. Thus, $\operatorname{rank}(P) = \operatorname{rank}(Q) = n$ requires both that the measurement windows distinguish all latent state directions and that the sampled trajectory represents them. This condition refers to the true latent dimension $n$; the measured Hankel rank alone cannot exclude additional unobserved or unexcited modes.

#### S1.1 State reconstruction from delay coordinates

When $\operatorname{rank}(P) = \operatorname{rank}(Q) = n$, equivalently $\operatorname{rank}(H_y) = n$, the latent state trajectory can be reconstructed from the delay data up to an invertible linear transformation. To obtain such a representation directly from the recordings, we use the compact singular value decomposition (SVD)

$$H_y = U_n \Sigma_n V_n^*. \tag{S2}$$

Here $U_n \in \mathbb{R}^{mp\times n}$ and $V_n \in \mathbb{R}^{T\times n}$ have orthonormal columns, $\Sigma_n \in \mathbb{R}^{n\times n}$ contains the $n$ nonzero singular values, and $*$ denotes conjugate transpose. All three matrices are computed from the recordings. Define $Z \in \mathbb{R}^{n\times T}$ and $\widehat{P} \in \mathbb{R}^{mp\times n}$ by

$$Z = U_n^* H_y = \Sigma_n V_n^*, \qquad \widehat{P} = U_n. \tag{S3}$$

Thus, $H_y = \widehat{P}Z$, and $\widehat{P}^\dagger = U_n^*$ maps each measured delay vector to its $n$-dimensional coordinates in $Z$.

These data-driven coordinates are related to the original latent states by an invertible linear transformation. Because $Q$ has full row rank, $H_y = PQ$ and $P$ have the same column space, so $P = U_n S$, where $S = U_n^* P \in \mathbb{R}^{n\times n}$ is invertible. Writing $X = Q \in \mathbb{R}^{n\times T}$ for the latent state sequence gives

$$Z = SX, \qquad \boldsymbol{z}_t = S\boldsymbol{x}_t. \tag{S4}$$

The reconstructed sequence therefore retains all information about the latent state in different coordinates. The computable map $\widehat{P}^\dagger$ generally differs from the theoretical inverse $P^\dagger$ in Eq. (6): it produces $\boldsymbol{z}_t$, rather than the original neural variables $\boldsymbol{x}_t$.

The recordings do not determine the particular choice of latent-state coordinates. For any invertible matrix $R \in \mathbb{R}^{n\times n}$, the factorisation $H_y = (PR^{-1})(RX)$ leaves the observations unchanged, while the state-evolution matrix in the transformed coordinates is $RAR^{-1}$. Additional information fixing the state coordinates is therefore required to identify the original $P$ and $\boldsymbol{x}_t$ themselves. The dynamical spectrum is unchanged because an invertible change of coordinates preserves eigenvalues.

##### S1.2 Reconstructing the dynamical spectrum from Hankel DMD

When $\operatorname{rank}(P) = \operatorname{rank}(Q) = n$, the Hankel DMD operator reproduces the latent dynamics on the span of the delay data. The Hankel DMD estimate $\tilde{A} = H_y' H_y^\dagger \in \mathbb{R}^{mp\times mp}$ in Eq. (17) is the minimum-Frobenius-norm minimiser of the least-squares problem

$$\min_{M\in\mathbb{R}^{mp\times mp}} \|H_y' - MH_y\|_F^2. \tag{S5}$$

Because $P$ has full column rank and $Q$ has full row rank, $(PQ)^\dagger = Q^\dagger P^\dagger$, $QQ^\dagger = I_n$, and $P^\dagger P = I_n$. Substituting $H_y = PQ$ and $H_y' = PAQ$ into $\tilde{A} = H_y' H_y^\dagger$ then gives

$$\tilde{A} = PAP^\dagger, \qquad \tilde{A}P = PA. \tag{S6}$$

Because $\operatorname{rank}(Q) = n$, $H_y = PQ$ and $P$ have the same column space. The identity $\tilde{A}P = PA$ therefore shows that applying $\tilde{A}$ to the delay-coordinate representation $P\boldsymbol{x}_t$ gives the same evolution as applying $A$ to the latent state and then mapping it through $P$.

The dynamical spectrum is obtained from the reduced Hankel DMD operator [1, 2]. Retaining $r$ singular directions gives

$$H_y \approx U_r \Sigma_r V_r^*, \qquad \tilde{A}_r = U_r^* H_y' V_r \Sigma_r^{-1}, \tag{S7}$$

where $U_r \in \mathbb{R}^{mp\times r}$ and $V_r \in \mathbb{R}^{T\times r}$ have orthonormal columns, $\Sigma_r \in \mathbb{R}^{r\times r}$ contains the retained singular values, and $\tilde{A}_r \in \mathbb{R}^{r\times r}$. When $\operatorname{rank}(P) = \operatorname{rank}(Q) = n$, we have $\operatorname{rank}(H_y) = n$, so retaining the full data rank gives $r = n$. With $S = U_n^* P \in \mathbb{R}^{n\times n}$, Eq. (S6) gives

$$\tilde{A}_n = SAS^{-1}. \tag{S8}$$

Thus, the reduced Hankel DMD operator has exactly the same eigenvalues as $A$, including their algebraic multiplicities, without requiring knowledge of the original state coordinates. These eigenvalues are the dynamical spectrum recovered from the recordings.

The unreduced Hankel DMD operator also contains the eigenvalues of $A$, together with additional zero eigenvalues associated with directions outside the span of the delay data. Because $P^\dagger$ vanishes on the orthogonal complement of $\mathrm{col}(P)$, the $mp-n$ additional directions contribute zero eigenvalues when $mp > n$. A genuine zero eigenvalue of $A$ is retained in the recovered spectrum and should not be confused with these additional zeros.

Equation (S8) is exact for the noise-free system and does not require $A$ to be invertible or diagonalisable. In noisy recordings, the retained rank $r$ must be estimated from the singular values, and weak or poorly conditioned directions can be difficult to recover accurately. If fewer than $n$ data directions are retained, Eq. (S8) no longer follows.

##### S1.3 Extension to nonlinear dynamics through Koopman theory

The same derivation applies to nonlinear dynamics when the measured observables belong to a finite-dimensional Koopman-invariant subspace. Consider the nonlinear system $\boldsymbol{x}_{t+1} = F(\boldsymbol{x}_t)$ with measurements $\boldsymbol{y}_t = h(\boldsymbol{x}_t)$ from Eq. (21), where $\mathcal{X} \subseteq \mathbb{R}^n$ is the state space, $F : \mathcal{X} \to \mathcal{X}$ is the state-evolution map, and $h : \mathcal{X} \to \mathbb{R}^p$ is the measurement function. Let $\mathcal{G}_d$ be a $d$-dimensional Koopman-invariant subspace containing the components of $h$, and let $\boldsymbol{\phi} = (\phi_1, \ldots, \phi_d)^\top : \mathcal{X} \to \mathbb{C}^d$ collect any basis of this subspace. There then exist matrices $K \in \mathbb{C}^{d\times d}$ and $C_K \in \mathbb{C}^{p\times d}$ such that

$$\boldsymbol{\phi}(F(\boldsymbol{x})) = K\boldsymbol{\phi}(\boldsymbol{x}), \qquad h(\boldsymbol{x}) = C_K\boldsymbol{\phi}(\boldsymbol{x}). \tag{S9}$$

Unlike the eigenfunction basis used in the main text, $\boldsymbol{\phi}$ may be any basis of the invariant subspace, so $K$ need not be diagonal or diagonalisable. With $\boldsymbol{w}_t = \boldsymbol{\phi}(\boldsymbol{x}_t) \in \mathbb{C}^d$, these relations give $\boldsymbol{w}_{t+1} = K\boldsymbol{w}_t$ and $\boldsymbol{y}_t = C_K\boldsymbol{w}_t$.

Using these coordinates, the Hankel matrices have the same factorised form as in the linear case. For delay depth $m$ and $T$ snapshot pairs, define $P_K \in \mathbb{C}^{mp\times d}$ and $Q_K \in \mathbb{C}^{d\times T}$ by

$$P_K = \begin{pmatrix} C_K \\ C_K K \\ \vdots \\ C_K K^{m-1} \end{pmatrix}, \qquad Q_K = \begin{pmatrix} \boldsymbol{w}_0 & K\boldsymbol{w}_0 & \cdots & K^{T-1}\boldsymbol{w}_0 \end{pmatrix}. \tag{S10}$$

The Hankel matrices satisfy

$$H_y = P_K Q_K, \qquad H_y' = P_K K Q_K, \qquad \mathrm{rank}(H_y) \le \min\{\mathrm{rank}(P_K), \mathrm{rank}(Q_K)\}. \tag{S11}$$

Thus, $P_K$ describes which directions in the invariant subspace are distinguishable through the delayed measurements, whereas $Q_K$ describes which directions are represented in the sampled trajectory.

When $\mathrm{rank}(P_K) = \mathrm{rank}(Q_K) = d$, the same reduced Hankel DMD construction recovers the eigenvalues of $K$. For noise-free data, let $H_y = U_d \Sigma_d V_d^*$ be the compact SVD, where $U_d \in \mathbb{C}^{mp\times d}$ and $V_d \in \mathbb{C}^{T\times d}$ have orthonormal columns and $\Sigma_d \in \mathbb{R}^{d\times d}$ contains the nonzero singular values. With $S_K = U_d^* P_K \in \mathbb{C}^{d\times d}$, which is invertible,

$$\tilde{A}_d = U_d^* H_y' V_d \Sigma_d^{-1} = S_K K S_K^{-1}. \tag{S12}$$

Therefore, the reduced Hankel DMD operator has the same eigenvalues as $K$, including their algebraic multiplicities. These are the Koopman eigenvalues represented in the selected invariant subspace. As in the linear case, the unreduced operator contains these eigenvalues together with additional zero eigenvalues associated with directions outside the span of the delay data.

When $K$ is diagonalisable, the representation in Eq. (S9) can be written in terms of Koopman eigenfunctions, as in the main text. Let $\boldsymbol{r}_i, \boldsymbol{\ell}_i \in \mathbb{C}^d$ be right and left eigenvectors of $K$ associated with $\lambda_i \in \mathbb{C}$ and normalised so that $\boldsymbol{\ell}_i^* \boldsymbol{r}_j = \delta_{ij}$. Then $\varphi_i : \mathcal{X} \to \mathbb{C}$ defined by

$$\varphi_i(\boldsymbol{x}) = \boldsymbol{\ell}_i^* \boldsymbol{\phi}(\boldsymbol{x}), \qquad (\mathcal{K}\varphi_i)(\boldsymbol{x}) = \lambda_i \varphi_i(\boldsymbol{x}) \tag{S13}$$

is a Koopman eigenfunction, and the measurements satisfy

$$\boldsymbol{y}_t = \sum_{i=1}^{d} (C_K \boldsymbol{r}_i) \lambda_i^t \varphi_i(\boldsymbol{x}_0). \tag{S14}$$

Choosing these eigenfunctions as coordinates gives the diagonal representation used in Eqs. (24)–(28) of the main text. If $K$ is not diagonalisable, Eqs. (S9) and (S12) still hold, although the mode expansion in Eq. (S14) must be replaced by the corresponding Jordan-chain representation.

#### S2 Simulations

This section specifies the simulations of Section 3.1 of the main text and extends them. Section S2.1 covers the linear oscillator network. It varies the stimulation site and the delay depth in the idealised feedforward setting of the main text, and repeats the analysis in a recurrent network with noise. Section S2.2 covers the Stuart–Landau oscillators.

##### S2.1 Linear simulation

###### S2.1.1 Network structure and observation

The linear network used in the simulation consists of 12 damped oscillators with $n = 24$ state variables and the system dynamics were sampled at $f_s = 500$ Hz (Table S1). Oscillator $k$ (denoted O$k$) was represented by the pair of state variables $(x_{2k-1}, x_{2k})$, which evolves by the rotation–scaling block of Methods 5.1 of the main text, which gives it the eigenvalue pair $\rho_k e^{\pm 2\pi i f_k / f_s}$. Five oscillators are slow-decaying ($\rho_k = 0.999$, a decay time constant of 2.0 s) and seven are fast-decaying ($\rho_k = 0.89$–0.96, 17–49 ms). Frequencies (4–95 Hz) are assigned independently of $\rho_k$. Only $p = 8$ of the 24 states are observed. The oscillators are coupled by 23 directed edges, all from a lower-indexed to a higher-indexed oscillator, so the network is strictly feedforward. An edge from oscillator $j$ to oscillator $k$ sets the off-diagonal block $A_{kj} = g_{kj}G$, where the gain $g_{kj} \in [0.12, 0.36]$ is listed in Table S1 and $G$ is one of two fixed $2 \times 2$ templates,

$$G_1 = \begin{pmatrix} 1.00 & 0.25 \\ -0.18 & 0.82 \end{pmatrix}, \qquad G_2 = \begin{pmatrix} 0.78 & -0.22 \\ 0.31 & 0.95 \end{pmatrix}.$$

Because every edge points downstream, $A$ is block lower triangular. Its eigenvalues are therefore exactly the 24 eigenvalues of the diagonal blocks, and an impulse into oscillator $j$ excites only the eigenvalue pairs of $j$ and of the oscillators downstream of it. The observation matrix $C$ selects both states of O1, O2, and O10 and the first state of O7 and O12, and with these channels the observability matrix $P_m$ reaches full rank at delay depth $m = 8$.

###### S2.1.2 Stimulation site and delay depth in the idealised feedforward setting

A stimulation site that reaches a larger portion of the network excites more eigenvalues, and a greater delay depth resolves more of the excited eigenvalues. Although the main text used a single site at the upstream end of the network (site i) and two delay depths, we additionally compare three sites with different downstream reach and sweep the delay depth, to show that the site sets a ceiling on recovery through rank $Q$ and that delay embedding cannot exceed it. The setting is that of Methods 5.1 of the main text. Coupling is strictly one-way and the network runs without noise, so each eigenvalue is either excited or has exactly zero amplitude, and every limit on recovery is an exact rank deficiency of $P_m$ or $Q$.

Runs and eigenvalue estimation follow Methods 5.1 of the main text, with three extensions.

**Table S1. Oscillators of the linear network.** ● marks an oscillator reached by the impulse at that site. ○ marks an oscillator that is not reached by the impulse but is present in the idealised feedforward setting, because it is slow-decaying and the initial state lies in the slow-decaying eigenspace. In that setting rank $Q$ is twice the number of marked rows. "Observed" gives the number of observed states of each oscillator, and the superscript on each gain gives the template, $G_1$ or $G_2$, of that edge.

| oscillator | $f_k$ (Hz) | $\rho_k$ | group | observed | outgoing edges (gain) | passive | i | ii | iii |
|---|---|---|---|---|---|---|---|---|---|
| O1 | 76 | 0.90 | fast | 2 | O2 $(.30)^1$, O3 $(.22)^2$, O5 $(.18)^1$ | | ● | | |
| O2 | 35 | 0.999 | slow | 2 | O4 $(.34)^2$, O5 $(.26)^1$, O7 $(.14)^2$ | ○ | ● | ○ | ○ |
| O3 | 54 | 0.92 | fast | – | O6 $(.32)^1$, O7 $(.24)^2$, O10 $(.12)^1$ | | ● | | |
| O4 | 83 | 0.95 | fast | – | O7 $(.30)^2$, O8 $(.20)^1$ | | ● | ● | |
| O5 | 95 | 0.999 | slow | – | O6 $(.28)^2$, O8 $(.32)^1$, O11 $(.16)^2$ | ○ | ● | ○ | ○ |
| O6 | 28 | 0.89 | fast | – | O9 $(.28)^1$, O11 $(.18)^2$ | | ● | | |
| O7 | 61 | 0.94 | fast | 1 | O9 $(.35)^1$, O10 $(.24)^2$ | | ● | ● | ● |
| O8 | 12 | 0.91 | fast | – | O10 $(.31)^1$, O11 $(.26)^2$ | | ● | ● | |
| O9 | 43 | 0.96 | fast | – | O12 $(.36)^1$ | | ● | ● | ● |
| O10 | 17 | 0.999 | slow | 2 | O12 $(.32)^2$ | ○ | ● | ● | ● |
| O11 | 9 | 0.999 | slow | – | O12 $(.27)^1$ | ○ | ● | ● | ○ |
| O12 | 4 | 0.999 | slow | 1 | – | ○ | ● | ● | ● |
| | | | | | rank $Q$ (idealised feedforward setting) | 10 | 24 | 18 | 14 |

First, the impulse is applied to the first state of O1 (site i), O4 (site ii), or O7 (site iii) (Fig. S1A). The impulse amplitude and the observation matrix are the same at every site, so differences in recovery arise from the stimulation site alone. Second, the delay depth is swept over $m = 1, \ldots, 20$. Third, rank $P_m$ and rank $Q$ are computed as numerical ranks with the same threshold as the SVD truncation, where $Q = [\boldsymbol{x}_0,\, A\boldsymbol{x}_0, \ldots, A^{79}\boldsymbol{x}_0]$ is built from the state $\boldsymbol{x}_0$ at the impulse. The recovery tolerance of 0.01 is less than one sixth of the smallest distance between two true eigenvalues (0.063), so a match cannot come from a neighbouring eigenvalue.

The initial state is constructed to contain only the slowly decaying eigenvalues. It is $2\,\mathrm{Re}\sum_j c_j \boldsymbol{v}_j$, summed over one eigenvector $\boldsymbol{v}_j$ of each slow-decaying conjugate pair, with $|c_j|$ drawn uniformly from $[0.8, 1.2]$ and a uniformly random phase. Because the slow-decaying eigenspace is invariant under $A$, the state stays in it until the impulse, and the fast-decaying eigenvalues enter the data only through the impulse.

In this setting rank $Q$ can be predicted by counting oscillators (Table S1). Because $A$ is diagonalisable with distinct eigenvalues, rank $Q$ equals the number of eigenvalues whose eigenvectors have a non-zero coefficient in the state at the start of the analysed trajectory. The initial state supplies the five slow-decaying pairs at every site, and the impulse adds the pairs of the oscillators downstream of the site. rank $Q$ is therefore twice the number of oscillators that are slow-decaying, downstream of the site, or both. Without an impulse this gives 10. Site i (O1) reaches all 12 oscillators, giving 24. Site ii (O4) reaches O4 and O7–O12, which with the slow-decaying O2 and O5 make nine, giving 18. Site iii (O7) reaches O7, O9, O10, and O12, which with O2, O5, and O11 make seven, giving 14.

The stimulation site set an exact ceiling on recovery, and delay embedding could not exceed it (Fig. S1). To isolate the effect of excitation, we first fixed the delay depth at $m = 8$, where rank $P_m = 24$ (Fig. S1B–E). Stimulation at the upstream site i propagated through the whole network, giving rank $Q = 24$ and recovery of all 24 eigenvalues. At site ii the impulse reached a smaller portion of the network, and rank $Q$ and recovery both fell to 18. At the further downstream site iii both fell to 14. The unrecovered eigenvalues were exactly those of the fast-decaying oscillators that the impulse did not reach, O1, O3, and O6 at site ii, and additionally O4 and O8 at site iii.

Sweeping the delay depth showed that each condition rose to its rank $Q$ and stayed there through $m = 20$ (Fig. S1F). Conditions with a smaller rank $Q$ reached their ceiling earlier, at $m = 4$ for the passive condition, $m = 6$ for sites ii and iii, and $m = 8$ for site i, because fewer excited directions had to be distinguished. Delay embedding therefore cannot recover eigenvalues that are absent from the trajectory.

##### S2.1.3 Recurrent noisy setting

The recurrent noisy setting examines whether the previous results still hold once the network is less idealised. Here, we remove two constraints. Weak reverse edges make the deterministic system reachable from every stimulation site, and a noise-driven baseline gives every eigenvalue non-zero amplitude almost surely. Recovery is limited instead by observability, excitation strength and the finite signal-to-noise ratio. If the stimulation site and delay depth still shape recovery here, their effects do not rely on the exact zeros of the idealised setting.

The first relaxation adds a weak reverse edge to every forward edge (Fig. S2A). The reverse edge from $k$ to $j$ sets $A_{jk} = \varepsilon\, g_{kj} G'$, where $G'$ is the template not used by the forward edge, and the diagonal blocks are left unchanged. We set $\varepsilon = 6.25 \times 10^{-4}$, small enough that the network stays stable (spectral radius 0.99938) and its eigenvalues stay close to those of the idealised setting. The true eigenvalues in this setting are computed from the recurrent $A$ itself.

The second relaxation adds process and observation noise,

$$\boldsymbol{x}_{t+1} = A\boldsymbol{x}_t + \sigma_w \boldsymbol{w}_t, \qquad \boldsymbol{y}_t = C\boldsymbol{x}_t + \sigma_v \boldsymbol{v}_t, \tag{S15}$$

where $\boldsymbol{w}_t \sim \mathcal{N}(\mathbf{0}, I_n)$ and $\boldsymbol{v}_t \sim \mathcal{N}(\mathbf{0}, I_p)$ are independent, with $\sigma_w = 10^{-3}$ and $\sigma_v = 10^{-5}$. The initial state is drawn from the stationary covariance of the noise-driven recurrent network and rescaled so that the observed activity has unit standard deviation. Noise makes the numerical rank full, so the SVD is truncated at $r = 24$, the true state dimension, and results are averaged over 10 noise realisations. The stimulation sites, impulse amplitude, and delay sweep are as in the idealised setting. For Fig. S2G, the impulse amplitude is additionally varied over $\alpha = 0$, 0.5, 1, 2, 3, 4, and 6 at $m = 20$, the depth of Fig. S2B–E.

The recurrent noisy setting reproduced the site ordering even though rank $Q = 24$ at every site (Fig. S2B–F). Averaged over 10 noise realisations at $m = 20$, the passive condition and sites iii, ii, and i recovered 10, 14, 18, and 21 of 24 eigenvalues (Fig. S2F). The first three match the idealised ceilings exactly. A representative realisation shows the same ordering, with 10, 14, 18, and 22 (Fig. S2B–E).

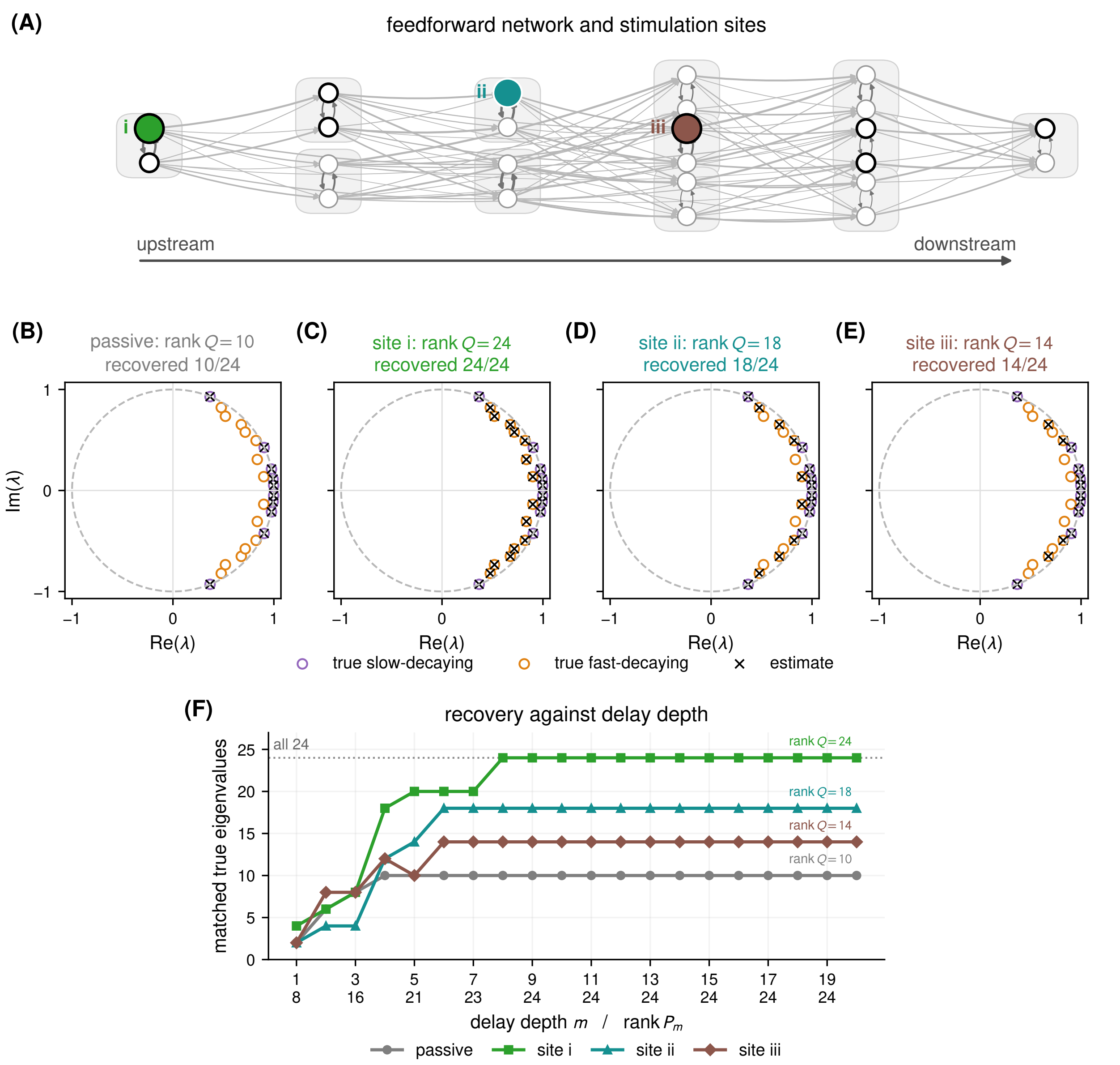

**Figure S1. In the idealised feedforward setting, the stimulation site sets an exact rank-$Q$ ceiling that delay embedding cannot exceed. (A)** The feedforward 24-state network. Grey boxes group the two states of each oscillator, black rings mark observed states, and coloured nodes identify the first state of O1, O4, and O7 stimulated at sites i, ii, and iii. **(B–E)** True eigenvalues (circles) and Hankel DMD estimates (crosses) at $m = 8$, where rank $P_m = 24$, for the passive condition and for stimulation at sites i, ii, and iii. Titles give rank $Q$ and the number of recovered eigenvalues, which are equal in every condition. **(F)** Recovered eigenvalues against delay depth. Annotations give rank $Q$, and the second row of tick labels gives rank $P_m$.

Delay embedding still raised recovery, but more gradually than in the idealised setting (Fig. S2F). rank $P_m$ was full from $m = 3$, yet recovery kept rising until about $m = 13$, whereas in the idealised setting it saturated by $m = 8$. The weak reverse edges make every direction observable in principle but many only weakly, so additional delays improved recovery without changing rank $P_m$, as expected from Section 2.4 of the main text. Recovery also increased with impulse amplitude and kept the site ordering at every nonzero amplitude (Fig. S2G), saturating from $\alpha \approx 2$ at 21, 18, and 14 eigenvalues for sites i, ii, and iii. Recovery tracked how strongly each mode entered the data. We weighted each mode by $\|C\boldsymbol{v}_i\|\,|\boldsymbol{w}_i^H \boldsymbol{x}_0|$, the product of its observability and its excitation, where $\boldsymbol{v}_i$ is the unit right eigenvector, $\boldsymbol{w}_i^H$ the matching row of the inverse eigenvector matrix, and $\boldsymbol{x}_0$ the state at the impulse. At $m = 20$, modes with a weight above $10^{-1.5}$ were recovered in 95% of cases and those below it in 17% (Spearman $\rho = 0.71$ across the 48 condition–mode-pair cells). Every mode had a nonzero weight, but the sites differed in how many they excited only weakly, below $10^{-1.5}$: none at site i, four at site ii, six at site iii, and eight in the passive condition. Of the 16 pairs left unrecovered across the four conditions, 15 were weakly excited. The exception was one pair at site i. The exact ceilings of the idealised setting thus became graded limits set by excitation strength, and the stimulation site still determined how many eigenvalues were recovered.

Thus, delay embedding, perturbation, and the stimulation site shaped recovery in much the same way once coupling was no longer strictly one-way and the network was noisy, even though rank $Q$ no longer distinguished the conditions. The site and delay-depth effects of the idealised setting therefore do not depend on its two idealisations.

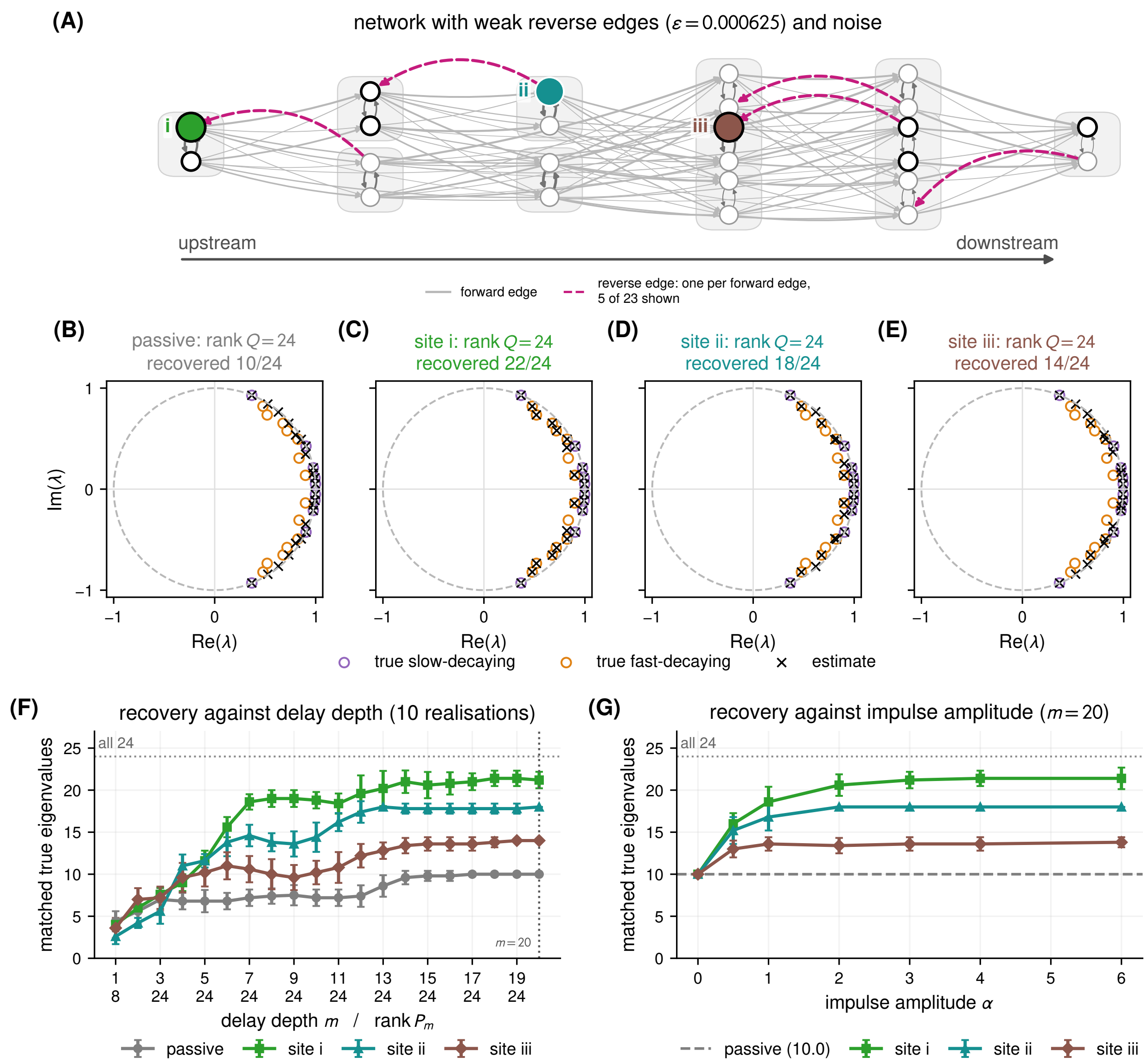

**Figure S2. In the recurrent noisy setting,** rank $Q$ **is full at every site, yet the stimulation site still orders recovery. (A)** The network with a weak reverse edge on every forward edge. Five of the 23 reverse edges are drawn as examples (dashed magenta). Process and observation noise are added. **(B–E)** True eigenvalues (circles) and Hankel DMD estimates (crosses) at $m = 20$ in a representative noise realisation whose counts follow the mean ordering, for the passive condition and for stimulation at sites i, ii, and iii. Titles give rank $Q$, now 24 in every condition, and the number of recovered eigenvalues. **(F)** Recovered eigenvalues against delay depth. Points are means over 10 noise realisations and bars their standard deviation. The dotted line marks $m = 20$, the depth of (B–E) and (G). **(G)** Recovery against impulse amplitude at $m = 20$. The dashed line is the passive condition.

##### S2.2 Nonlinear simulation with Stuart–Landau oscillators

This section derives the Koopman eigenvalues targeted in the Stuart–Landau simulation (Section 3.1.2 and Methods 5.2), gives its full numerical specification, and shows that its result holds across impulse sizes and observation matrices. The model and the names of the target eigenvalues follow Section 3.1.2.

###### S2.2.1 Koopman eigenvalues in linear observations

The eigenvalues follow from two Koopman eigenfunctions of each oscillator. The squared amplitude obeys

$$\frac{d|a_j|^2}{dt} = 2r_j|a_j|^2 - 2|a_j|^4, \tag{S16}$$

so the radial function $\varphi_j = 1 - r_j/|a_j|^2$ evolves as $\varphi_j(t) = \varphi_j(0)\, e^{-2r_j t}$. It is therefore a Koopman eigenfunction with eigenvalue $-2r_j$, and it vanishes on the limit cycle. The phase $\arg a_j$ advances at the amplitude-dependent rate $\gamma_j - \beta_j|a_j|^2$. Correcting it for this dependence gives the phase function

$$\Theta_j = \arg a_j + \frac{\beta_j}{2} \log \frac{r_j}{|a_j|^2}, \tag{S17}$$

which advances uniformly as $\Theta_j(t) = \Theta_j(0) + \omega_j t$. Thus $e^{i\Theta_j}$ is a Koopman eigenfunction with eigenvalue $i\omega_j$.

Inverting these two relations expresses the complex amplitude in terms of the eigenfunctions,

$$a_j = \sqrt{r_j}\, e^{i\Theta_j}\, (1 - \varphi_j)^{-(1+i\beta_j)/2}. \tag{S18}$$

When $|a_j|^2 > r_j/2$, so that $|\varphi_j| < 1$, a binomial expansion yields

$$a_j(t) = \sum_{k=0}^{\infty} d_{j,k}\, e^{(-2kr_j + i\omega_j)t}, \qquad d_{j,k} = \sqrt{r_j}\, c_{j,k}\, \varphi_j(0)^k\, e^{i\Theta_j(0)}, \tag{S19}$$

where $c_{j,k}$ are the binomial coefficients of $(1 - \varphi_j)^{-(1+i\beta_j)/2}$. This is the expansion used in the main text. On the limit cycle, $\varphi_j(0) = 0$ and only the $k = 0$ term remains. Near the cycle, the coefficient of order $k$ scales as $\varphi_j(0)^k$, so higher orders contribute progressively less and require larger displacements to be expressed.

The full Koopman spectrum also contains the harmonics in $\omega_j$ and their combinations across oscillators, but these do not appear in linear observations. The system is invariant under rotation of each $a_j$, so the expansion of $a_j$ contains only eigenfunctions of the form $\varphi_j^k e^{i\Theta_j}$. The oscillators are uncoupled, so $a_j$ contains no eigenfunctions of the other oscillators. Because the observations are real, each term appears together with its complex conjugate. The eigenvalues expressed in $\boldsymbol{y} = C\boldsymbol{x}$ are therefore exactly the non-decaying eigenvalues $\pm i\omega_j$ and the fast-decaying eigenvalues $-2kr_j \pm i\omega_j$.

**Table S2. Stuart–Landau parameters and recovery across impulse sizes. (A)** Model parameters and the real parts of the fast-decaying eigenvalues of orders $k = 1$ and 2. **(B)** Mean number of oscillators (out of 4) whose non-decaying and order-$k$ fast-decaying eigenvalues are recovered at $m = 200$, averaged over five observation matrices. "Kept" is the number of DMD eigenvalues remaining after the amplitude and stability filters.

**(A) Model parameters**

| $j$ | $f_j$ (Hz) | $\omega_j$ (rad s$^{-1}$) | $r_j$ (s$^{-1}$) | $\beta_j$ | $\sqrt{r_j}$ | $-2r_j$, $-4r_j$ (s$^{-1}$) |
|---|---|---|---|---|---|---|
| 1 | 4 | 25.1 | 0.6 | 0.6 | 0.77 | −1.2, −2.4 |
| 2 | 8 | 50.3 | 0.8 | −0.4 | 0.89 | −1.6, −3.2 |
| 3 | 15 | 94.2 | 1.2 | 0.8 | 1.10 | −2.4, −4.8 |
| 4 | 30 | 188.5 | 2.0 | −0.7 | 1.41 | −4.0, −8.0 |

**(B) Recovery against impulse size**

| $\alpha$ | non-decaying | $k = 1$ | $k = 2$ | $k = 3$ | kept |
|---|---|---|---|---|---|
| 0 | 4.0 | 0.0 | 0.0 | 0.0 | 8.0 |
| 0.05 | 4.0 | 3.8 | 0.0 | 0.0 | 15.6 |
| 0.1 | 4.0 | 4.0 | 0.6 | 0.0 | 17.2 |
| 0.2 | 4.0 | 4.0 | 3.2 | 0.2 | 24.4 |
| 0.3 | 4.0 | 4.0 | 3.4 | 1.0 | 28.0 |
| 0.5 | 3.8 | 3.8 | 3.2 | 2.0 | 32.0 |
| 0.8 | 4.0 | 4.0 | 3.2 | 1.8 | 40.4 |

###### S2.2.2 Simulation and estimation

Table S2A lists the parameters. Trajectories are computed from the closed-form solution of Eq. (30), so they carry no integration error. They are sampled every $\Delta t = 2$ ms for 4 s (2000 samples). In the passive condition, each oscillator starts on its limit cycle at a random phase. In the perturbed condition, the impulse direction $\boldsymbol{b}$ is the all-ones vector, so every one of the eight real state coordinates moves by $\alpha$ and each amplitude shifts as $a_j \to a_j + \alpha(1 + i)$. The observation matrix $C \in \mathbb{R}^{2\times 8}$ has independent Gaussian entries and unit-norm rows. The main-text figure uses one draw of $C$, and Table S2B averages over five.

Eigenvalues are estimated with DMD ($m = 1$) and Hankel DMD ($m = 200$), with the SVD rank chosen by the optimal hard threshold [3]. Discrete-time estimates $\mu$ with $|\mu| \geq 1.05$ or with amplitude below 1% of the largest are discarded, and the rest are converted to continuous time as $\lambda = \log \mu/\Delta t$. As in the linear simulation, target and estimated eigenvalues are paired one-to-one, so each estimate can recover at most one target. A target eigenvalue counts as recovered when its paired estimate lies within $0.4 \times 2r_j$ of its real part and within 7.5 rad s$^{-1}$ of its imaginary part. The real-part window is 0.4 times the spacing $2r_j$ between successive orders, and the imaginary-part window is 0.3 times the smallest spacing between the $\pm\omega_j$, so an estimate cannot be credited to a neighbouring order or frequency.

##### S2.2.3 Robustness across impulse sizes and observation matrices

The main-text result held across impulse sizes and observation matrices (Table S2B). In Fig. 2F, no target eigenvalue was recovered at $m = 1$. At $m = 200$, the four non-decaying eigenvalue pairs were recovered in both conditions, and the fast-decaying eigenvalues appeared only with the impulse, for all four oscillators at $k = 1$ and three at $k = 2$. Averaged over five observation matrices at $m = 200$, the non-decaying eigenvalues were recovered at every impulse size. Without the impulse, exactly eight estimates survived the filters, and these were the four non-decaying pairs, as expected from the absence of harmonics in linear observations. The fast-decaying eigenvalues required the impulse, and higher orders required larger displacements, consistent with the scaling of $d_{j,k}$ with $\varphi_j(0)^k$. The $k = 1$ eigenvalues were recovered for nearly all four oscillators already at $\alpha = 0.05$. The $k = 2$ eigenvalues began to appear at $\alpha = 0.1$ and were recovered for 3.2 of 4 oscillators at the main-text value $\alpha = 0.2$. The $k = 3$ eigenvalues were recovered for at most one oscillator on average below $\alpha = 0.5$.

### S3 Analysis of the ECoG data

This section supplements the ECoG analysis of Section 3.2 of the main text. It describes how stimulation sites were selected (Section S3.1), how eigenvalue estimates were validated (Section S3.2), how the post-stimulus change appears without the frequency and decay thresholds (Section S3.3), and how robust it is to the rank and validation criteria (Section S3.4).

#### S3.1 Inclusion of participant–site pairs and artefact removal

Stimulation sites entered the analysis in three stages (Table S3A). The first stage kept participant–site pairs whose stimulation protocol met the criteria of Methods 5.3 of the main text, giving 1788 candidate pairs from 50 participants. The second stage screened channels and trials with the checks in Table S3B, and 1503 pairs retained at least ten passing trials. The third stage required every fit at a site, with both methods and in both windows, to reach rank 50, so that counts of validated estimates are compared at a common rank (Methods 5.4 of the main text). This removed 185 sites and six participants and left the *primary dataset* of 1318 stimulation sites from 44 participants, which is used throughout unless stated otherwise.

The quality checks remove amplifier artefacts before they enter the analysed windows. A raw-epoch check scans each trial for plateaus and persistent offsets over −310 to −10 ms and +10 to +900 ms. It runs on the unprocessed epoch, before window-wise mean removal, because plateaus and offsets are absolute-level effects that centring hides, and its thresholds are scaled by each channel's pre-stimulus variability. A window check then rejects any analysis window with non-finite samples or a flat channel. A channel that fails the raw-epoch check in many trials is removed from the pair, and a trial is removed when any remaining channel fails. The recording channels are the good ECoG channels of the dataset, excluding the two stimulated contacts.

**Table S3. Selection of stimulation sites, attrition under multi-step decay validation, and quality-control rules. (A)** Site selection and attrition of candidate eigenpairs. Candidate eigenpairs are counted over single-trial fits and are not distinct physiological processes. For candidate eigenpairs, "residual" is the number that passed the one-step residual criterion and "multi-step" the number that also passed multi-step decay validation. **(B)** Channel and trial quality checks. The pre-stimulus scale $s$ is 1.4826 times the median absolute deviation over −310 to −10 ms (at least 1 $\mu$V), and $\tilde{x}$ is the pre-stimulus median.

**(A) Site selection and attrition under multi-step decay validation**

| level | condition | before | after | removed |
|---|---|---|---|---|
| sites | stimulation criteria → channel and trial QC | 1788 | 1503 | 285 |
| sites | → all fits reach rank 50 | 1503 | 1318 | 185 |
| participants | → all fits reach rank 50 | 50 | 44 | 6 |
| | | residual | multi-step | removed |
| candidate eigenpairs | DMD, pre-stimulus | 111,983 | 68,265 | 39.0% |
| | DMD, post-stimulus | 125,626 | 73,080 | 41.8% |
| | Hankel, pre-stimulus | 99,748 | 96,119 | 3.6% |
| | Hankel, post-stimulus | 120,119 | 112,766 | 6.1% |

**(B) Channel and trial quality checks**

| check | rule |
|---|---|
| plateau | Over +10 to +900 ms, consecutive samples whose first difference is at most 1 $\mu$V while $x$ stays far from $\tilde{x}$. A channel is flagged when such a run lasts at least 20 ms with $\lvert x - \tilde{x}\rvert \geq 20\,s$, or at least 50 ms with $\lvert x - \tilde{x}\rvert \geq 10\,s$. |
| offset | $\Delta$ is the median over +300 to +900 ms minus $\tilde{x}$. A channel is flagged when $\lvert\Delta\rvert > 20\,s$ and $\lvert\Delta\rvert > 1000$ $\mu$V, or when $\lvert\Delta\rvert > 2000$ $\mu$V and the signal stays on the same side at least $\lvert\Delta\rvert/2$ away from $\tilde{x}$ for at least 300 ms. |
| channel exclusion | A channel is removed from the pair when it is flagged in at least $\max(2, \lceil N/2 \rceil)$ of the $N$ candidate trials. At least four channels must remain. |
| trial exclusion | A trial is removed when any remaining channel is flagged, or when either analysis window has a channel with fewer than 99.9% finite samples or a standard deviation below $10^{-10}$. |
| trial selection | Ten trials are taken, evenly spaced in time, from those that pass. Pairs with fewer than ten are dropped. |

##### S3.2 Held-out validation of eigenvalue estimates

Candidate eigenpairs are validated as in Methods 5.5 of the main text. Each of the ten trials of a site serves once as the training trial, and a candidate eigenpair fitted to it is validated when it is stable with a frequency between 0 and 100 Hz, its one-step held-out residual (main-text Eq. (31)) is at most 0.1 in at least five of the nine held-out trials, and it passes multi-step decay validation. This section explains why the multi-step check is needed and how it behaves.

The one-step residual cannot reliably detect an overstated decay rate. Over a single sample, a fitted decay that is too fast changes the predicted eigenfunction only slightly, so such an estimate can pass the residual criterion. Multi-step decay validation instead compares the fitted decay $|\lambda_j|^\ell$ over a lag of $\ell$ samples, the number nearest to 5 ms, with the decay $|\hat{\mu}_j^{(q)}|$ that the eigenfunction values show over the same lag on held-out trial $q$ (main-text Eq. (32)). A candidate eigenpair is excluded when

$$\text{median}_q\, |\hat{\mu}_j^{(q)}| - |\lambda_j|^\ell > 0.05.$$

The rule is one-sided and is applied identically to both methods. It removes estimates whose fitted decay is faster than the held-out data support, which is the error that would inflate the count of fast-decaying eigenvalues. The check therefore works against the main-text finding that perturbation increases this count, not in its favour.

Figure S3A–D shows the two criteria applied in sequence at an example stimulation site. Grey points fail the residual criterion, magenta points pass it but fail multi-step decay validation, and teal points pass both. Most magenta points come from standard DMD and lie far inside the unit circle, where a short fitted time constant is most likely to be overstated.

The rule rarely rejected a correct decay and almost always rejected an overstated one (Fig. S3E,F). On synthetic 20-Hz eigenfunctions, it rejected the true decay in at most 2.8% of nine-trial sets across the evaluable settings, but rejected a fourfold-overstated decay in at least 99% of noise-free transients and stationary processes with $\tau \leq 50$ ms. In the ECoG data it removed mainly short-time-constant candidates from standard DMD (Fig. S3G), 39.0% and 41.8% of the residual-passing candidate eigenpairs in the pre- and post-stimulus windows, compared with 3.6% and 6.1% for Hankel DMD (Table S3A).

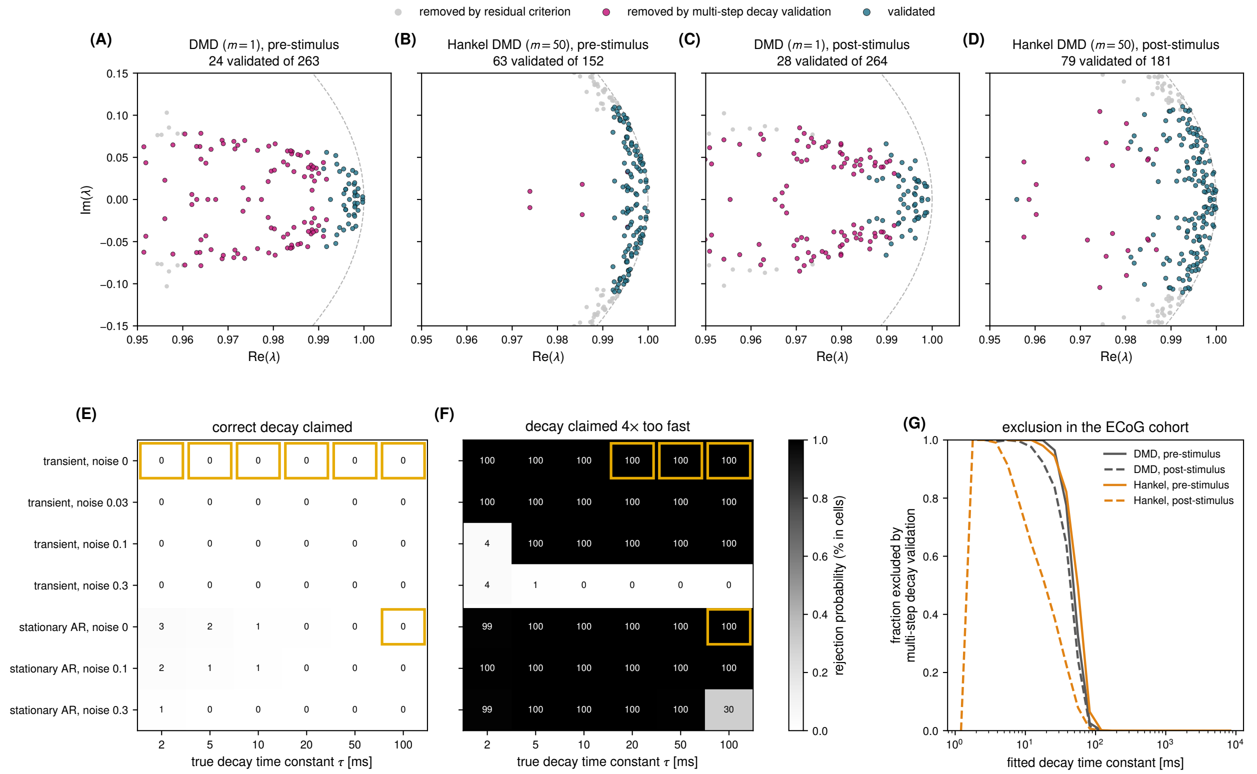

**Figure S3. Selection of candidate eigenpairs and the behaviour of multi-step decay validation.** **(A–D)** Candidate eigenpairs on the eigenvalue plane at an example stimulation site: pre-stimulus (A,B) and post-stimulus (C,D) windows, from DMD ($m = 1$) and Hankel DMD ($m = 50$). Every point is a stable 0–100-Hz candidate eigenpair from one of ten single-trial fits. Grey points fail the one-step held-out residual criterion, magenta points pass the residual criterion but fail 5-ms multi-step decay validation, and teal points pass both criteria and are validated. The plane shows both members of each conjugate pair, whereas title counts use frequencies $f \geq 0$. **(E,F)** Synthetic calibration of the 5-ms rule with tolerance 0.05 on 20-Hz eigenfunctions with known decay. Cells give the percentage of 500 nine-trial sets rejected when the candidate eigenpair claims the true decay (E) or a decay four times faster (F). Gold frames mark settings in which some sets pass the residual criterion. **(G)** Fraction of residual-passing ECoG candidate eigenpairs rejected by multi-step decay validation, against fitted decay time constant. The rule is one-sided: it removes estimates whose fitted decay is faster than the held-out data support and does not test whether slower fitted decay is accurate.

##### S3.3 Post-stimulus change in the frequency–decay distribution

Without the thresholds of main-text Fig. 3, the post-stimulus change appears as a shift of Hankel DMD estimates from long to short time constants (Fig. S4). After stimulation, Hankel DMD yielded fewer validated estimates at time constants of 0.3–3 s and more at approximately 20–250 ms (Fig. S4B). The gain was concentrated between 5 and 40 Hz, with almost none above 40 Hz. Standard DMD showed a much weaker gain, at time constants below 100 ms and frequencies below 30 Hz (Fig. S4A). The map is a difference between two distributions, so the loss at long time constants and the gain at short ones need not involve the same estimates.

Neither threshold of the main text drives this result. The gain peaks near 120 ms, close to the 100-ms time constant ($-10\ \mathrm{s}^{-1}$) that separates fast- from slow-decaying eigenvalues (Fig. S4D). Moving this threshold would change the number of validated estimates of fast-decaying eigenvalues but not the shift itself. Most of the gain also lies above 8 Hz (Fig. S4C), consistent with the frequency split of main-text Fig. 3F.

The position of the peak also explains why the slow-decaying count barely changed after stimulation (main-text Fig. 3E). Because the peak lies just above the threshold, part of the gain is counted as slow-decaying. With Hankel DMD, the gain between 100 and 215 ms (+1.29 estimates/site/fit) was largely offset by losses at longer time constants within the map (−1.48 in total, −1.19 of it between 0.3 and 3.2 s). The net change on the slow-decaying side of the threshold within the map was therefore only −0.19, against +1.63 on the fast-decaying side. Estimates with time constants beyond the 10-s display range contributed a further −0.13, giving a total slow-decaying change of −0.32.

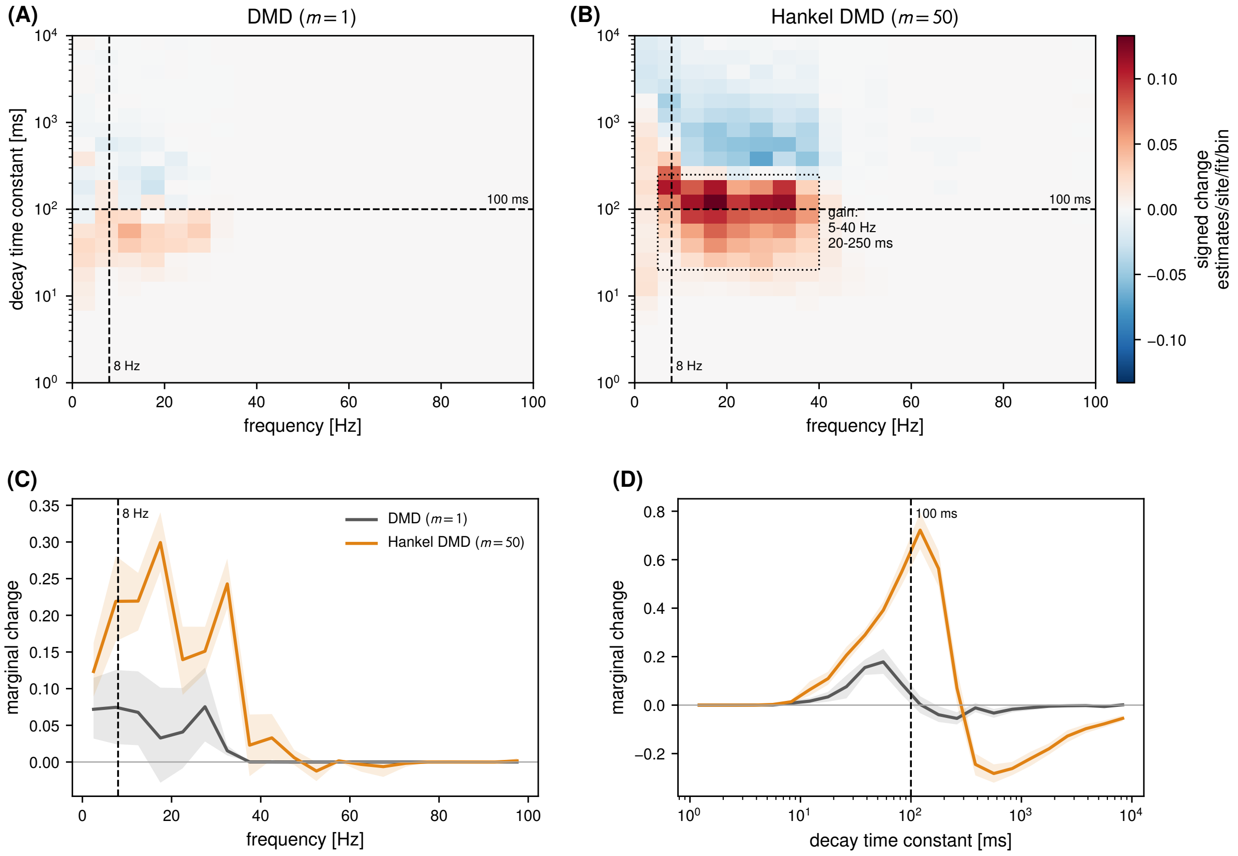

**Figure S4. After stimulation, Hankel DMD yields fewer estimates with time constants of 0.3–3 s and more with time constants of 20–250 ms at 5–40 Hz.** Primary dataset (1318 sites from 44 participants). **(A,B)** Differences between the post- and pre-stimulus windows in the number of validated estimates, for standard and Hankel DMD. Bins are 5 Hz by one-sixth decade over 1–10,000 ms. Values are normalised per site and fit within participant, then averaged over 44 participants. Dashed lines are the two thresholds of main-text Fig. 3, at 8 Hz and at a 100-ms time constant ($-10$ s$^{-1}$). The dotted box in (B) outlines the Hankel DMD gain at 5–40 Hz and 20–250 ms. (A) and (B) share one colour scale. **(C,D)** Frequency and decay-time marginals, the sums of the bin-wise changes over decay time and over frequency, with the same thresholds. Shading is a descriptive 95% participant-bootstrap interval.

#### S3.4 Sensitivity to rank and validation criteria

The post-stimulus increase in validated estimates of fast-decaying eigenvalues with Hankel DMD remained positive across all ranks and validation criteria examined (Fig. S5). Under every setting examined, it was positive in every participant and exceeded the increase with standard DMD.

The increase was unchanged when the rank was lowered, whether or not the lower rank also admitted more sites (Fig. S5A). In the primary dataset at rank 50, the mean increase was 1.627 estimates/site/fit with Hankel DMD (median 1.547, positive in all 44 participants), against 0.556 with standard DMD. The median corresponds to the within-participant increase of 15.5 per site in the main text, which pools the ten fits. Refitting the same 1318 sites from 44 participants at rank 30, which changes the rank alone, gave 1.657 with Hankel DMD (positive in all 44 participants) and 0.801 with standard DMD (43 of 44). The rank-50 requirement also excluded 185 sites (Section S3.1). At rank 30 these sites can be included, and the full set of 1503 quality-controlled sites from 50 participants gave 1.612 and 0.687. With Hankel DMD the increase was positive in all 50 participants, and with standard DMD in 45.

The residual criterion changed the size of the Hankel DMD increase but not its sign (Fig. S5B). Across thresholds of 0.05, 0.1, and 0.2 and requirements of 3, 5, and 7 of 9 held-out trials, the mean increase ranged from 0.79 to 2.86 and was positive in all 44 participants.

Multi-step decay validation mattered for standard DMD but hardly for Hankel DMD (Fig. S5C). Across validation horizons of 2, 5, and 10 ms and tolerances of 0.05 and 0.10, the increase was 1.63–1.87 with Hankel DMD and 0.56–1.00 with standard DMD. Without multi-step decay validation it was 1.89 and 1.32. Removing overstated decays thus cut the standard DMD increase by more than half but reduced the Hankel DMD increase by less than 15%, in line with the attrition in Table S3A.

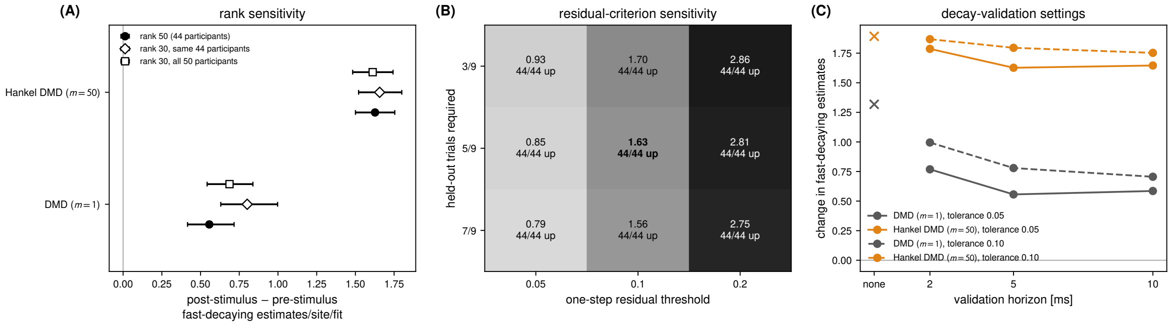

**Figure S5. The post-stimulus increase in validated estimates of fast-decaying eigenvalues from Hankel DMD is robust to rank and validation settings.** Values are post-stimulus minus pre-stimulus differences in estimates per site per fit, averaged over participants with each participant weighted equally. **(A)** Primary dataset at rank 50 (filled circles), the same 1318 sites and 44 participants refitted at rank 30 (diamonds), and all 1503 sites from 50 participants at rank 30 (squares). Bars are descriptive 95% participant-bootstrap intervals. **(B)** Hankel DMD across one-step residual thresholds and required held-out trials. Cells give the mean change and the number of participants with an increase, and bold marks the primary criterion. **(C)** Both methods without multi-step decay validation (crosses) and across validation horizons and tolerances.
